# Evolutionary arms race between SARS-CoV-2 and interferon signaling via dynamic interaction with autophagy

**DOI:** 10.1101/2023.11.13.566859

**Authors:** Jae Seung Lee, Mark Dittmar, Jesse Miller, Minghua Li, Kasirajan Ayyanathan, Max Ferretti, Jesse Hulahan, Kanupriya Whig, Zienab Etwebi, Trevor Griesman, David C. Schultz, Sara Cherry

**Affiliations:** Department of Pathology and Laboratory Medicine, University of Pennsylvania, Philadelphia; PA 19104, USA; Department of Microbiology, University of Pennsylvania, Philadelphia; PA 19104, USA; Department of Biochemistry and Biophysics, University of Pennsylvania; Philadelphia, PA 19104, USA

**Author notes:** Department of Pathology, University of Texas Medical Branch, Galveston, TX 77555, USA. Corresponding author. (C.S.).

**Keywords:** SARS-CoV-2, Alpha and Omicron variants, Interferon response, Innate immune, Viral evolution, Autophagy, ATG7, ATG9A, OPTN, MAVS, ORF9b

## Abstract

SARS-CoV-2 emerged, and is evolving to efficiently infect humans worldwide. SARS-CoV-2 evades early innate recognition, interferon signaling activated only in bystander cells. This balance of innate activation and viral evasion has important consequences, but the pathways involved are incompletely understood. Here we find that autophagy genes regulate innate immune signaling, impacting the basal set point of interferons, and thus permissivity to infection. Mechanistically, autophagy genes negatively regulate MAVS, and this low basal level of MAVS is efficiently antagonized by SARS-CoV-2 ORF9b, blocking interferon activation in infected cells. However, upon loss of autophagy increased MAVS overcomes ORF9b-mediated antagonism suppressing infection. This has led to the evolution of SARS-CoV-2 variants to express higher levels of ORF9b, allowing SARS-CoV-2 to replicate under conditions of increased MAVS signaling. Altogether, we find a critical role of autophagy in the regulation of innate immunity and uncover an evolutionary trajectory of SARS-CoV-2 ORF9b to overcome host defenses.

## Introduction

Severe acute respiratory syndrome coronavirus 2 (SARS-CoV-2) was identified in Wuhan China in December 2019, and caused a global pandemic ^1,2^. SARS-CoV-2 initially infects the respiratory epithelium ^3^, and innate immune recognition by the epithelial barrier is the first line of defense against viruses including coronaviruses ^4^. Cytoplasmic sensors including retinoic acid-inducible gene-I (RIG-I) like receptors (RLRs) bind viral RNAs and are activated to engage the mitochondrial adapter protein MAVS to induce downstream signaling ^5,6^. The oligomerization of MAVS at mitochondria ultimately leads to IRF3 phosphorylation and translocation to the nucleus and the induction of interferons (IFNs) ^6^. Secreted IFNs activate the JAK-STAT pathway which induces the expression of interferon stimulated genes (ISGs) which block viral infection ^4,7^. Decreased signaling leads to poor outcomes ^8–10^, and increased IFN signaling prior to infection can be protective ^11,12^. Therefore, a better understanding of how IFN is regulated may both allow better prediction of outcomes and be harnessed for new interventions.

Autophagy is a catabolic cellular mechanism that captures cytoplasmic components for degradation and maintains homeostasis regulating diverse pathways including stress responses and innate immunity ^13–15^. Autophagy is a stepwise process that involves many conserved genes ^16^. Diverse stimuli and stressors activate the Unc-51-like kinases (ULK1/2) which initiates the formation of a vesicle nucleation complex including VPS34 (PIK3C3). After initiation, autophagy elongation factors including ATG7 process ATG8 family proteins (ATG8s/LC3s) into lipidated forms that localize to the growing autophagosome. To elongate the phagophore, additional membrane source material is supplied by ATG9A containing vesicles derived from various cellular organelles ^17^. Cargo adapters including p62 (SQSTM1) and OPTN bring specific substrates to the growing autophagosome ^18^. Indeed, mitochondria can be selectively captured by a process called mitophagy and recent studies have demonstrated an important role for ATG9A and OPTN in this process ^19–21^. Next, the fully formed autophagosome fuses with the lysosome leading to the degradation of captured cargo such as mitochondria during mitophagy ^15^.

Autophagy can impact viral infections in diverse ways including directly through the regulation of replication or indirectly by regulating host processes including innate immunity ^13,22–24^. Previous studies on autophagy during SARS-CoV-2 infection have found pro-viral roles for autophagy in non-respiratory cells ^24–28^. Studies in Vero and Huh7 cells, found that some early autophagy factors (VPS34, DFPC1, and DFPC2) promote the establishment of replication organelle, while components involved in downstream steps in the pathway including autophagy elongation factors are dispensable or restrict infection of SARS-CoV-2 ^24,25^. Furthermore, ectopic expression studies in Vero, and Hela cells have shown that SARS-CoV-2 ORF3a, and ORF7a can block auto-lysosomal fusion; however, it is unclear if SARS-CoV-2 impacts this step during infection ^27,29,30^.

Previous studies have shown that autophagy can indirectly impact some viral infections by regulating innate immune signaling ^22,31,32^. In some contexts, immune signaling factors including RIG-I, STING, MDA5 and MAVS can be perturbed by the autophagy machinery ^22,33–38^. This dynamic change has high potential to affect the viruses sensitive to IFN treatment. However, it is unknown if autophagy controls innate immune signaling in respiratory cells and how this may impact SARS-CoV-2 infection. Indeed, SARS-CoV-2 evades early innate immune recognition in epithelial cells, leading to delayed interferon responses and increased pathogenesis ^10,39–44^. Thus, SARS-CoV-2-dependent induction of IFNs occurs largely in uninfected bystander cells at late time points post infection ^4,12,45^. Nevertheless, while SARS-CoV-2 can evade early recognition, this can be overcome by exogenous treatments that induce IFNs and IFN signaling ^12,46^. Many mechanisms have been proposed to explain how SARS-CoV-2 evades recognition ^10,39–43^; however, most studies used ectopic expression systems leading to the question of which viral proteins antagonize during bona fide infection ^12,39,41,42,45,47–52^. Moreover, many studies of SARS-CoV-2 use cell lines engineered with entry factors to promote infection (e.g. A549-ACE2) and are poorly innate immune responsive (e.g. HEK293T, HeLa) making it difficult to uncover how these signaling pathways impact replication in their endogenous contexts ^12,52,53^. In contrast, the human respiratory Calu-3 cells endogenously express entry factors, are permissive to infection, and are innate immune responsive, inducing and responding to viral PAMPs, Type I, and Type III IFNs ^12,45,52^. This system allows us to uncover the pathways that regulate SARS-CoV-2 infection including innate immune signaling.

As SARS-CoV-2 has continued to spread within human populations, new waves of variants continue to arise ^54^. Much of this evolution is in the Spike protein to evade pre-existing antibodies from vaccination or previous infections ^54–57^. Less is known about the evolutionary trajectory of other SARS-CoV-2 proteins ^44^. In this study we found that autophagy promotes SARS-CoV-2 infection. Using genetic approaches, we found that diverse components involved in autophagy as well as mitophagy negatively regulate the mitochondrial signaling adapter MAVS required for IFN induction. Loss of autophagy factors led to increased MAVS and IFNs which block infection. Indeed, we can restore infection by blocking IFN signaling in autophagy-deficient cells. Moreover, loss of autophagy promotes IFN induction not only in bystander cells but also in infected cells. Autophagy-dependent regulation of MAVS maintains MAVS at low levels that can be antagonized by the viral accessory protein ORF9b. However, ORF9b is expressed at low levels such that loss of autophagy leads to increased MAVS that overcomes ORF9b-mediated attenuation. Furthermore, we found that SARS-CoV-2 variants (Alpha, and Omicron) express higher level of ORF9b than the ancestral variant, which can overcome the increased levels of MAVS when autophagy is disabled. Therefore, these emerging variants can antagonize IFN activation more efficiently. Altogether, these data suggest that that SARS-CoV-2 variants are evolving to be less sensitive to innate immune defenses.

## Results

### Autophagy promotes SARS-CoV-2 replication in human respiratory epithelial cells

Autophagy is a complex cell biological process, and VPS34 (PIK3C3) is a key enzyme involved in the induction of autophagy ^15^. We and others found that treatment with the VPS34 inhibitor, VPS34 IN1, blocks SARS-CoV-2 infection ^15,58–60^. Moreover, previous study suggested that autophagy promotes viral replication factories ^24^. Therefore, we set out to define the role of autophagy in infection of the respiratory tract. We validated that treatment with the VSP34 inhibitor reduced infection and found >1000-fold reduction in human respiratory Calu-3 cells as measured by RT-qPCR monitoring either subgenomic N mRNA or genomic RNA (nsp14) (Fig 1a, and Extended Data Fig. 1a). Furthermore, SARS-CoV-2 protein levels were reduced by VPS43 IN1 as measured by immunoblot against nucleocapsid (N), and ORF7a (Extended Data Fig. 1b). We complemented these studies using three more differentiated human airway models. First, treatment of primary human bronchial epithelial cells in air liquid interface (ALI) with VPS34 IN1 reduced infection by >100-fold (Fig. 1b, and Extended Data Fig. 1c). Second, we used primary human nasal epithelial ALI cultures, and again observed significant reduction in infection upon VPS34 IN1 treatment (Extended Data Fig.1d). Lastly, we tested human iPSC-derived alveolar epithelial type 2 cells (iAT2) and found that VPS34 IN1 also inhibited infection in these cells (Extended Data Fig. 1e).

**Fig. 1.**
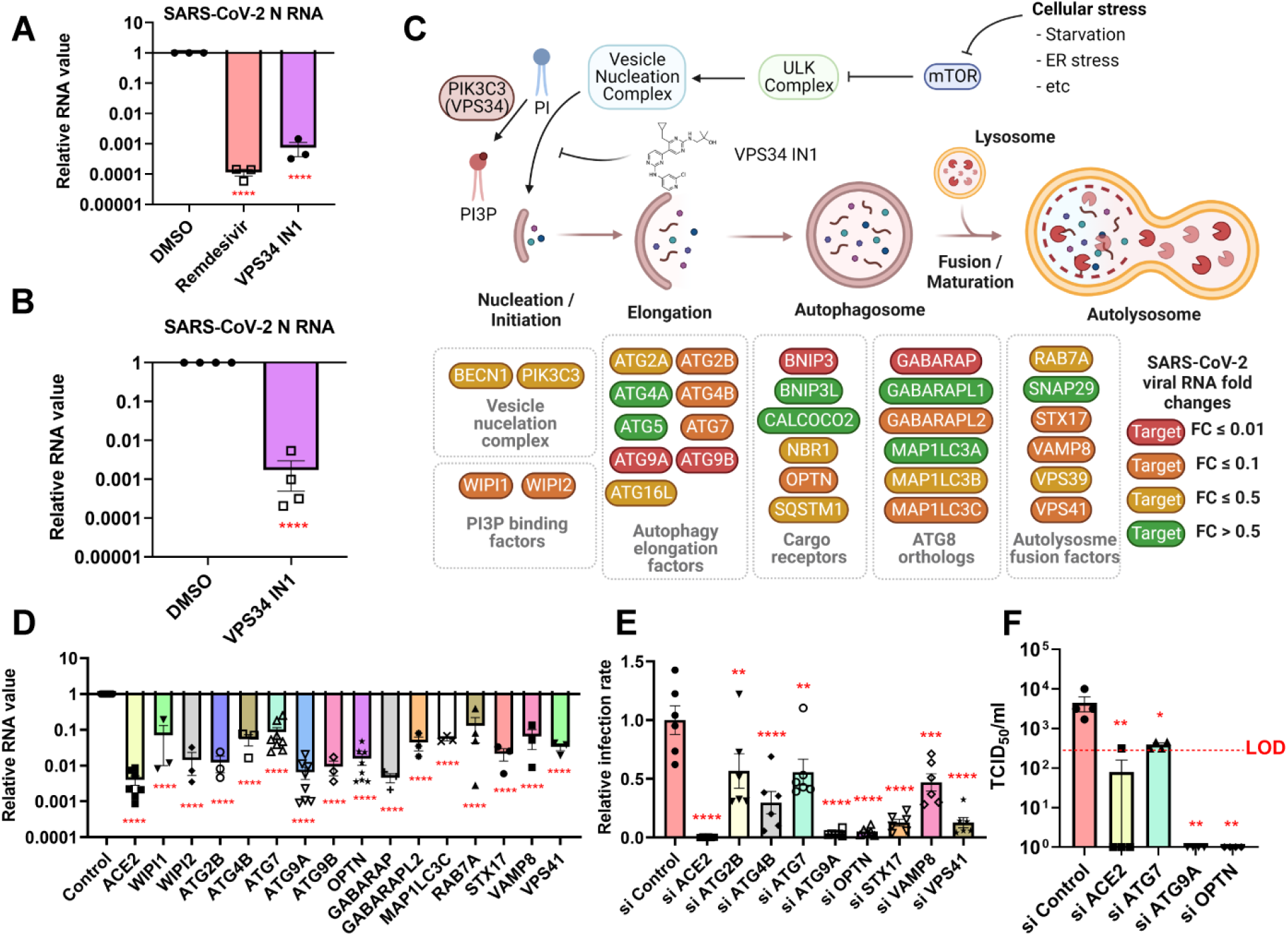
Autophagy promotes SARS-CoV-2 replication in human respiratory epithelial cells. (**a**) Calu-3 cells were pretreated with vehicle (DMSO) or 10 μM of the indicated drugs and infected with SARS-CoV-2 (MOI of 0.2) for 48h and total RNA was subject to RT-qPCR analysis of viral infection (N mRNA). (**b**) Human bronchial air-liquid interface cells were pretreated with vehicle or 3 μM VPS34 IN1 and infected with SARS-CoV-2 (MOI of 0.2). RT-qPCR analysis of viral infection (N mRNA) at 72 hpi. (**c**) Schematic of autophagy pathway and the impact of depletion of each gene in Calu-3 cells on SARS-CoV-2 replication as measured by RT-qPCR. With fold change of viral RNA levels compared to siRNA control (siControl). (**d** and **e**) Calu-3 cells were transfected with the indicated siRNAs and at 48 hr infected with SARS-CoV-2 (MOI of 0.2) for 48h and subject to (d) RT-qPCR analysis of viral RNA (N mRNA) compared to siControl or (e) processed for automated microscopy and automated image analysis quantified total cell numbers (nuclei+) and the percent infection (Spike+/nuclei+). (**f**) Calu-3 cells were transfected with the indicated siRNAs. 2 days post transfection, the cells were infected with SARS-CoV-2 (MOI of 0.2) and media replaced 4 hpi. At 48 hpi, the supernatant was titered and the TCID50s shown. (LOD, limit of detection). For all graphs, data shown are means ± SEMs with individual biological replicates shown. Significance was calculated using one-way ANOVA for all figures (*p < 0.05; ** p < 0.01; *** p < 0.001; **** p < 0.0001). ns, no significance.

Given the role of VPS34 in the initiation of autophagy, we next set out to genetically test the role of autophagy genes in SARS-CoV-2 infection of respiratory Calu-3 cells that are amenable to genetic manipulation, ^12,52,53^. Previous studies on autophagy during SARS-CoV-2 infection have focused on non-respiratory cells and thus we tested the role of these factors in Calu-3 cells ^24–28^. We used siRNAs to deplete 31 diverse components of the autophagy pathway involved in distinct steps from initiation to autophagolysosomal fusion (Fig. 1c, Extended Data Fig. 2a and 2b). As a control, we found that depletion of the entry receptor, ACE2, led to ∼ 100-fold reduction in infection as measured by RT-qPCR (Fig. 1d). We found that depletion of diverse autophagy factors from each step in the pathway including WIPI1, WIPI2, ATG2B, ATG4B, ATG7, ATG9A, ATG9B, BNIP3, OPTN, GABARAP, GABARAPL2, MAP1LC3C, RAB7A, STX17, VAMP8, and VPS41 decreased SARS-CoV-2 RNA >10-fold (Fig. 1c, and d). We complemented these studies by depleting each gene and monitoring infection by a microscopy-based assay where we used automated imaging and image analysis to quantify both cell viability and the percent infection (Fig. 1e, Extended Data Fig. 2c, and 2d). Again, we used ACE2 as a control, and found >100-fold reduction in infection demonstrating concordance between these assays. We observed striking reductions in infection upon depletion of autophagy genes ATG7, OPTN and ATG9A in both assays, and importantly depletion of these factors had no impact on cell viability (Extended Data Fig. 2c). Furthermore, we monitored the impact of ATG7, ATG9A, and OPTN on viral titers. As a control, we found that depletion of ACE2 led to >100-fold reduction in titers (Fig. 1f). Depletion of ATG9A or OPTN lead to >1000-fold reduction to below the limit of detection while depletion of ATG7 led to ∼ 10-fold reduction in viral titers (Fig. 1f). Altogether, we found that diverse autophagy genes promote SARS-CoV-2 infection and in particular, factors including that are involved in selective processes including mitophagy.

### Autophagy genes control SARS-CoV-2 infection through interferon signaling

Next, we set out to determine how autophagy may promote infection. Since the entry factors ACE2 and TMPRSS2 are essential for viral infection, we tested whether autophagy regulates their expression and found that loss of autophagy genes did not lead to significant reductions in mRNA levels of these factors (Extended Data Fig. 2e). Autophagy can regulate replication directly or indirectly through diverse pathways including innate immune signaling ^31,61,62^. Thus, we explored the interactions between autophagy genes and the interferon pathway. We focused these mechanistic studies on ATG7, OPTN and ATG9A since they are involved in disparate steps in the autophagy pathway and significantly impact infection without affecting cell viability (Fig. 1c to f, and Extended Data Fig. 2c). Moreover, OPTN and ATG9A promote selective mitophagy suggesting that they may regulate a specific signaling event at mitochondria.

We compared control-depleted, ACE2-depleted and cells depleted for the autophagy genes ATG7, ATG9A or OPTN and infected these cells with SARS-CoV-2. We verified that depletion of ACE2, or autophagy factors led to significant decreases in infection as measured by RT-qPCR (Fig. 2a, and Extended Data Fig. 3a). Furthermore, as expected, we found that SARS-CoV-2 infection of control cells led to increased expression of Type I and Type III IFNs, IFNB and IFNL1, as well as the interferon stimulated gene (ISG), TRIM22 at 48h post infection (Extended Data Fig. 3b to 3d). Moreover, as expected, depletion of the entry receptor ACE2 both blocked infection and led to reduced interferon signaling (Fig. 2a to g). This is consistent with a block to infection prior to innate immune activation. In contrast, and unexpectedly, loss of autophagy genes blocked infection but led to increased levels of IFNB, IFNL1, and the ISG TRIM22 (Fig. 2b to g, and Extended Data Fig. 3a).

**Fig. 2.**
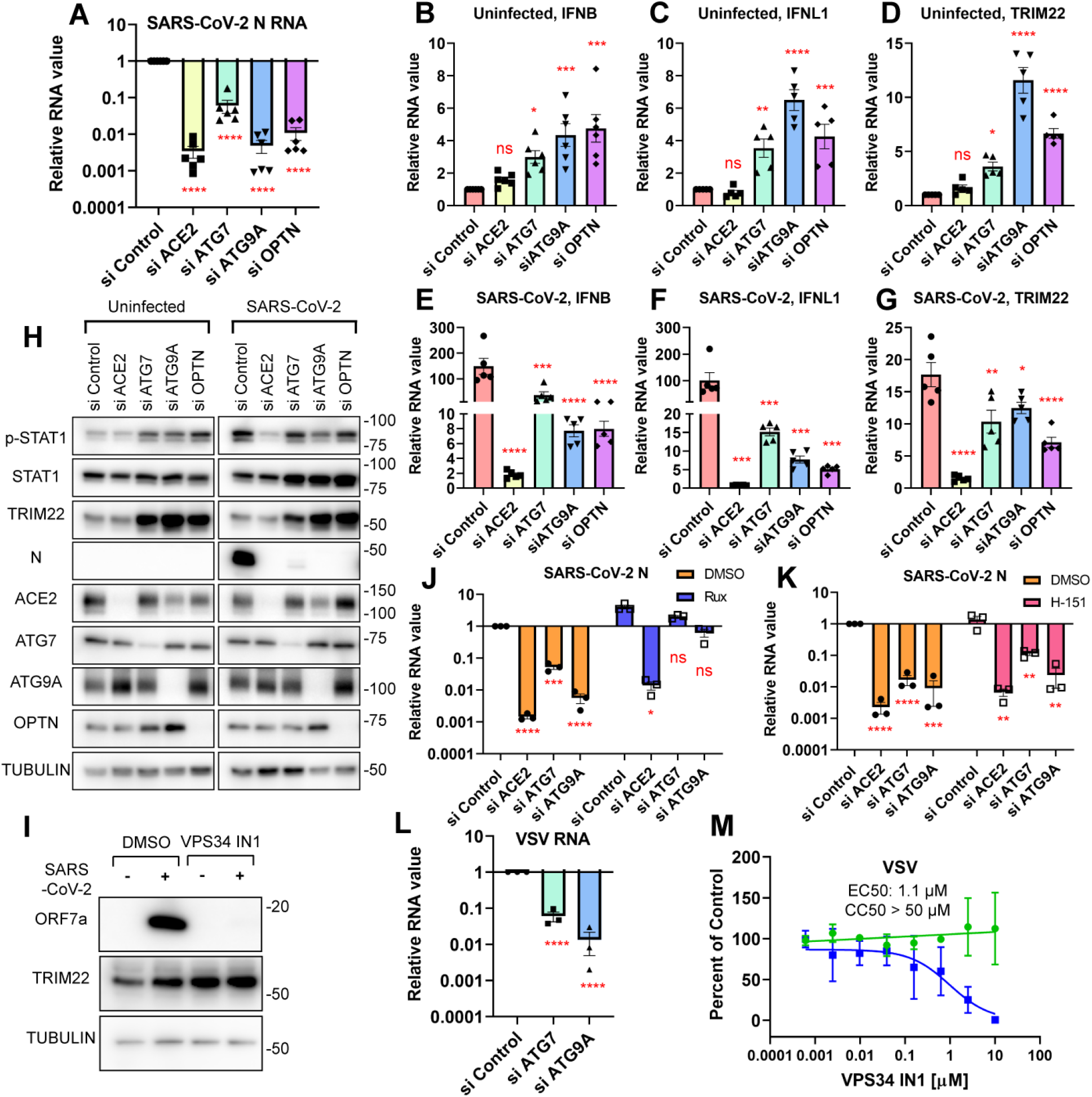
Autophagy controls SARS-CoV-2 infection through interferon signaling. (**a** to **h**) Calu-3 cells were transfected with the indicated siRNAs, at 48 h cells were either uninfected or infected with SARS-CoV-2 (MOI of 0.2) as indicated. (a to g) At 48 hpi, total RNA was subjected to RT-qPCR analysis of the indicated gene compared to siControl (h) total protein analyzed by immunoblot with the indicated antibodies. Representative blots shown for n=3. (**i**) Calu-3 cells were pretreated with vehicle (DMSO) or 2 μM VPS34 IN1, followed by infection with SARS-CoV-2 (MOI of 0.2). At 48 hpi, total protein was collected for immunoblot. Representative blot shown for n=3. (**j** and **k**) Calu-3 cells were transfected with the indicated siRNAs, after 2 days, cells were treated with vehicle (DMSO), 10 μM Ruxolitinib or 10 μM H-151 as indicated followed by infection with SARS-CoV-2 (MOI of 0.2). At 48 hpi viral RNA levels were analyzed by RT-qPCR (N mRNA) and normalized to vehicle treated siControl sample. (**l**) Calu-3 cells were transfected with the indicated siRNAs, 48 h post transfection cells were infected with VSV-GFP (MOI of 0.1). At 24 hpi, total RNA was subject to RT-qPCR for viral RNA (N region). (**m**) Calu-3 cells were pre-treated with vehicle (DMSO) or the indicated concentrations of VPS34 IN1 and infected with VSV-GFP (MOI of 1.8). At 24 hpi, the cells were processed for automated imaging and automated image analysis quantified total cell numbers (green, nuclei+) and the percent infection (blue, GFP+/nuclei+) shown as percent of vehicle control. For all graphs, data shown are means ± SEMs from indicated replicates. Significance was calculated using one-way ANOVA for (a to g, and l) or two-way ANOVA for (j and k) (* p < 0.05; ** p < 0.01; *** p < 0.001; **** p < 0.0001). ns, no significance.

We next determined if increased IFN mRNA and ISG mRNA production observed upon loss of autophagy genes also led to increased ISG protein expression. Indeed, loss of autophagy genes leads to increased TRIM22 in the absence of infection as measured by immunoblot (Fig. 2h). We also monitored the activation of status of the IFN signaling pathway. We monitored STAT1 activation and found increased levels of phospho-STAT1 upon depletion of autophagy genes in the absence of infection (Fig. 2h). Treatment with the VPS34 inhibitor, as with genetic depletion, also led to increased TRIM22 expression (Fig. 2i). These results demonstrate that autophagy negatively regulates interferon signaling suggesting that this may be the mechanism by which autophagy controls SARS-CoV-2 infection.

If autophagy impacts infection only through interferon signaling, we should be able to restore infection in autophagy-depleted cells by blocking interferon signaling and downstream antiviral ISG production. In contrast, if autophagy directly regulates viral replication, treatment with an inhibitor of interferon signaling should have little impact on infection of cells lacking autophagy. We used the JAK/STAT inhibitor Ruxolitinib and found that treatment of control cells led to a modest increase in infection accompanied by a striking reduction in the expression of the ISG TRIM22 (Fig. 2j, and Extended Data Fig. 3e to 3h) ^12,63^. In contrast, we found that viral infection was dramatically restored in ATG7 and ATG9A depleted cells upon Ruxolitinib treatment (Fig. 2j, and Extended Data Fig. 3e to 3g). These data show that autophagy genes control infection indirectly through the regulation of interferon signaling. Moreover, these data also suggest that autophagy is not required for SARS-CoV-2 replication.

Interferons can be induced by a number of upstream signaling pathways including STING ^4^. Previous studies have shown that STING is regulated by autophagy including ATG9A ^33^; and that STING activation can block SARS-CoV-2 infection ^12^. Therefore, we tested whether STING was involved in this autophagy-dependent pathway. We tested whether the STING inhibitor H-151 could restore infection in autophagy-depleted cells as we found with the JAK inhibitor ^33,35^. In contrast to Ruxolitinib treatment, H-151 had no impact on SARS-CoV-2 infection in control, or autophagy depleted cells (Fig. 2k and Extended Data Fig. 3i to 3k) ^12^. This demonstrates that autophagy-dependent control of SARS-CoV-2 infection is dependent on interferon signaling, independent of STING.

Given that diverse RNA viruses are controlled by IFNs, our data suggests that autophagy-dependent regulation of interferons may also impact infection of additional viruses that are sensitive to IFNs, such as vesicular stomatitis virus (VSV) (Extended Data Fig. 3l) ^64^. Indeed, we found that VSV replication is significantly decreased in ATG7 or ATG9A depleted cells (Fig. 2l). Moreover, VSV infection is inhibited by the VPS34 inhibitor with IC50 similar to SARS-CoV-2 (0.78 µM, Fig. 2m) ^59^. Thus, autophagy-dependent regulation of interferons impacts viral infections more broadly in respiratory epithelial cells.

### Autophagy controls IRF3 translocation in infected cells

Interferons are classically induced by RLRs downstream of RNA virus infections, including coronaviruses ^5^. We set out to explore how autophagy genes impact interferon activation basally and found that depletion of ATG7 or ATG9A enhanced the expression of both the Type I interferon IFNB, and type III interferon IFNL1 (Fig. 3b and c). Next, we used the canonical RLR stimulator Sendai virus (SeV) and found as expected, SeV induces IFNs rapidly, and depletion of either ATG7 or ATG9A potentiated SeV-induced activation of these IFNs (Fig. 3d and e). Likewise, VPS34 inhibitor treatment increased the expression of IFNB and IFNL1 basally, and upon SeV treatment (Fig. 3g to j).

**Fig. 3.**
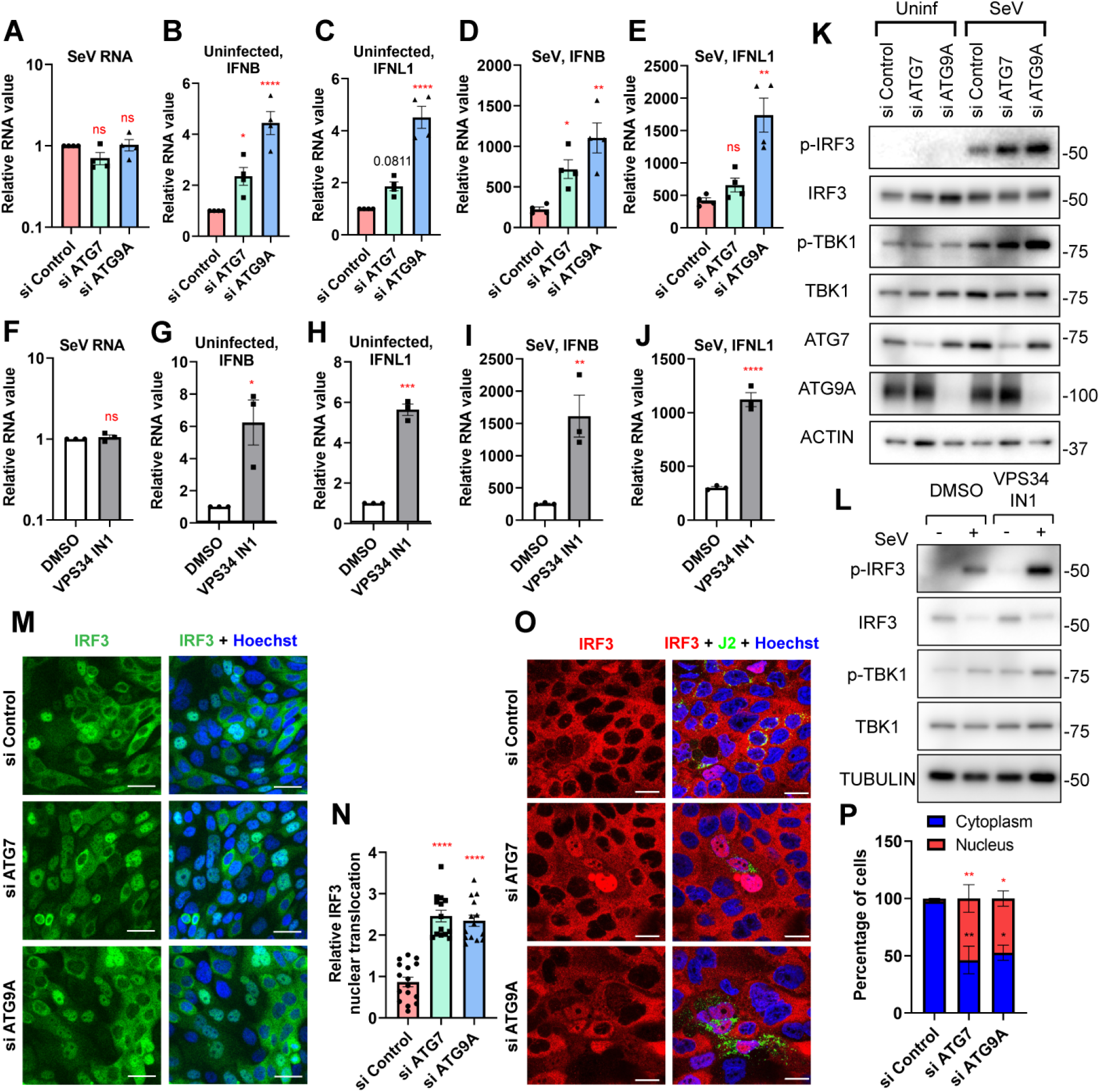
Autophagy blocks IRF3 translocation in virus infected cells. (**a** to **e**) Calu-3 cells were transfected with the indicated siRNAs, and 3 days later, either uninfected or infected with Sendai virus (SeV, 100 HAU/ml) for 6 hr. Total RNA was analyzed by RT-qPCR for the indicated targets. (**f** to **j**) Calu-3 cells were pretreated with vehicle (DMSO) or 2 μM VPS34 IN1 and uninfected or infected with SeV for 6 hr. Total RNA was analyzed by RT-qPCR for the indicated targets. (**k**) Calu-3 cells were transfected with the indicated siRNAs, 3 days later either uninfected or infected with SeV (100 HAU/ml) for 2 hr and protein lysates were subject to immunoblot analysis. Representative blot of n=3 shown. (**l**) Calu-3 cells were pretreated with vehicle (DMSO) or 2 μM VPS34 IN1 and uninfected or infected with SeV (100 HAU/ml) for 2 hr and protein lysates were analyzed by immunoblot with the indicated antibodies. Representative blot of n=3 shown. (**m** and **n**) Calu-3 cells were transfected with the indicated siRNAs, 3 days post transfection, infected with SeV (100 HAU/ml) for 6 hr. The cells were fixed and processed for confocal microscopy (anti-IRF3, green; nuclei, Hoechst 33342, blue) and nuclear IRF3 translocation was quantified. The scale bar represents 40 μm. (**o** and **p**) Calu-3 cells were transfected with the indicated siRNAs, 2 days post transfection, infected with SARS-CoV-2 for 48 hr. The cells were fixed and processed for confocal microscopy (anti-IRF3, red; anti-dsRNA, green; nuclei, Hoechst 33342, blue). Infection and nuclear IRF3 translocation were quantified. ATG depletion increased IRF3 nuclear localization in infected cells. The scale bar represents 20 μm. For all graphs, data shown are means ± SEMs from biological replicates. Significance was calculated using one-way ANOVA for (a to e, n, and p) or unpaired t-test for (f to j) (* p < 0.05; ** p < 0.01; *** p < 0.001; **** p < 0.0001). ns, no significance.

Since we observed increased IFN expression upon loss of autophagy genes and upon SeV treatment, these data suggest that autophagy functions within this pathway. RLR activation leads to phosphorylation and activation of TBK1 which phosphorylates the transcription factor IRF3 that subsequently translocates from the cytoplasm to the nucleus to induce expression of IFN genes ^5^. Thus, we explored the levels and activation status of components of this RLR pathway first by immunoblot. As expected, SeV activated the RLR pathway as measured by increased phosphorylation of TBK1 and IRF3 (Fig. 3k and l). Moreover, loss of autophagy genes or VPS34 inhibition led to further increases in phospho-TBK1 and phospho-IRF3. Second, we complemented these studies with confocal microscopy to explore how autophagy genes impact IRF3 nuclear translocation at single cell resolution. As expected, SeV treatment leads to robust translocation of IRF3 (Extended Data Fig. 4a). Upon depletion of autophagy genes, or treatment with the VPS34 inhibitor, we observed significant increased IRF3 translocation (Fig. 3m, n, and Extended Data Fig. 4b and 4c).

### Loss of autophagy genes leads to cell autonomous activation of IRF3 in SARS-CoV-2 infected cells

SARS-CoV-2 evades early detection by RLRs in infected cells, and IRF3 translocation is observed only in bystander cells at late time points post infection ^12,45^. We next determined if autophagy genes play a role in this process. As expected, IRF3 translocation was induced by SARS-CoV-2, but only uninfected bystander cells (Fig. 3o, and p). However, upon depletion of autophagy genes, ATG7 or ATG9A, IRF3 nuclear localization is now readily observed in SARS-CoV-2 infected cells (Fig. 3o, and p). Therefore, our data demonstrate that the autophagy pathway regulates signaling upstream of IRF3 during SARS-CoV-2 infection.

### Autophagy genes negatively regulate the mitochondrial signaling adapter MAVS

Since we found that autophagy negatively regulates TBK1 and IRF3, we next explored the upstream components of the signaling pathway. Viral RNA recognition by the cytoplasmic RLRs, RIG-I and MDA5, activates these receptors to bind MAVS at mitochondria, which leads to TBK1 activation ^6,65,66^. Moreover, the expression of these proteins impacts signaling: increased expression of RIG-I, MDA5 or MAVS activates IRF3 and IFN expression ^67^. Thus, we hypothesized that the expression level of one of these signaling proteins is regulated by autophagy to control IFN.

We first monitored the protein expression levels of RIG-I, MDA5 and MAVS upon depletion of autophagy genes ATG7 or ATG9A, and observed increased expression of all three proteins (Fig. 4a). Since many proteins within the IFN signaling pathway are themselves ISGs, we also determined the dependence on IFN signaling. Thus, we treated autophagy-depleted cells with Ruxolitinib, and found that the increased MDA5 and RIG-I expression was reduced by Ruxolitinib. We also monitored the mRNA levels of these factors, and found that MDA5 and RIG-I are indeed ISGs (Fig. 4b to c). In contrast, we found that MAVS protein and mRNA expression are not controlled by IFN signaling; the autophagy-dependent increase in MAVS protein levels is IFN-independent suggesting that autophagy directly impact MAVS (Fig. 4a and d).

**Fig. 4.**
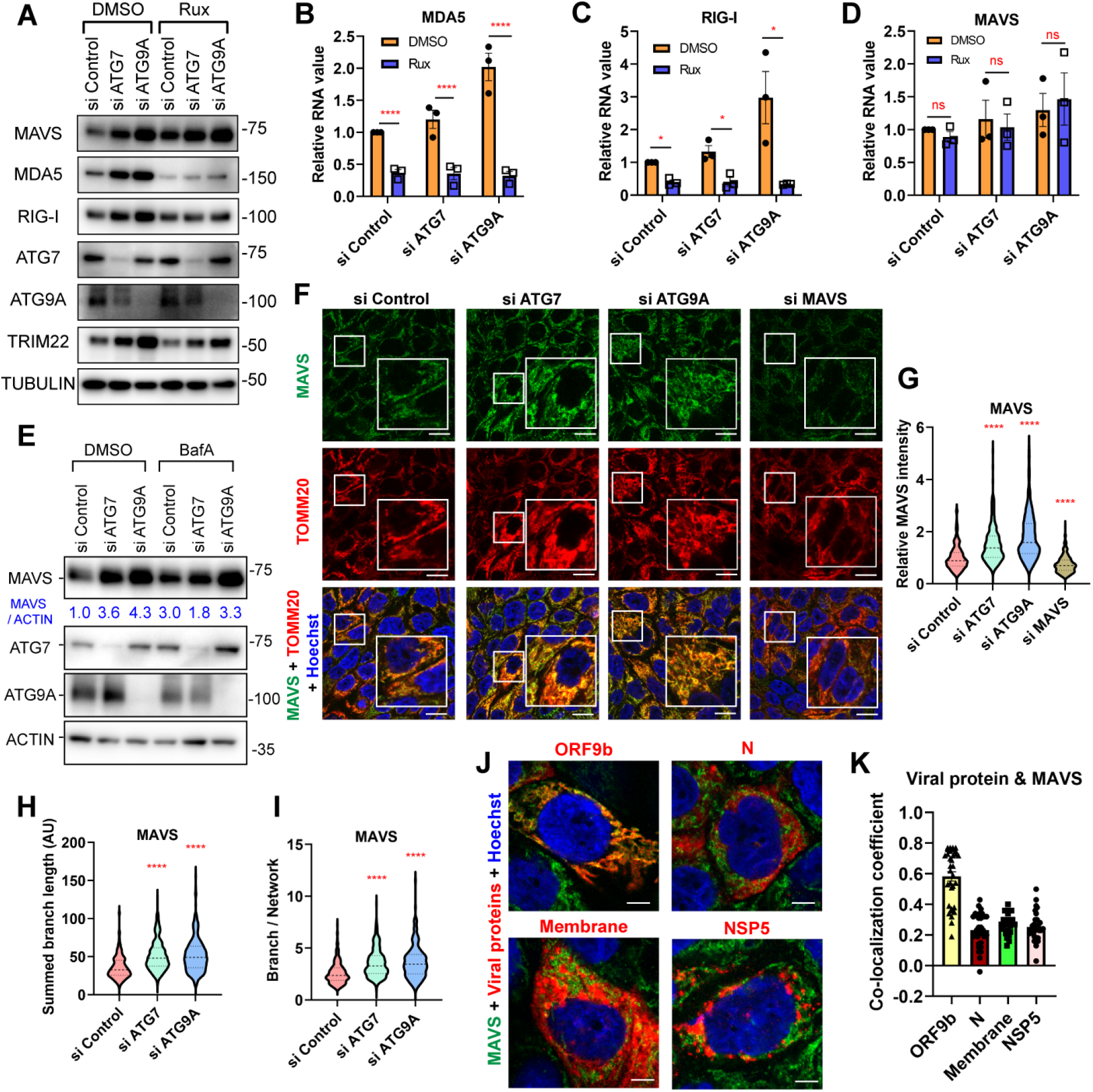
Autophagy genes regulate interferon signaling pathway via MAVS. (**a** to **d**) Calu-3 cells were transfected with the indicated siRNAs, 2 days post transfection, either treated with vehicle (DMSO) or 10 μM Ruxolitinib for 24 hr. (a) Total protein was collected for immunoblot with the indicated antibodies. Representative blot of n=3 shown. (b to d) Total RNA was collected for RT-qPCR with the indicated target. (**e**) Calu-3 cells were transfected with the indicated siRNAs for 3 days. Vehicle (DMSO) or 0.1 μM BafilomycinA1 was added for 16 hr, and protein samples were collected for immunoblot with the indicated antibodies. MAVS intensity normalized to ACTIN is shown. Representative blot of n=3 shown. (**f** to **i**) Calu-3 cells were transfected with the indicated siRNAs for 3 days and processed for confocal microscopy (anti-MAVS, green; mitochondria, anti-TOMM20, red; nuclei, Hoechst 33342, blue). The scale bar represents 20 μm. Large square is magnified image from inset. (g to i) More than 250 cells for each condition were analyzed across 3 independent experiments. (g) MAVS signal intensity was measured, MAVS morphology was analyzed by measuring (h) length and (i) number of branches in a network. (**j** and **k**) Calu-3 cells were infected with SARS-CoV-2 (MOI of 0.2) for 24 hr and processed for microscopy (anti-MAVS, green; anti-ORF9b, anti-nucleocapsid (N), anti-membrane, or anti-NSP5, red; nuclei, Hoechst 33342, blue). The scale bar represents 5 μm. More than 25 cells were analyzed to measure co-localization co-efficient across three independent experiments and each cell shown as individual dot. For all graphs, data shown are means ± SEMs from biological replicates. Significance was calculated using one-way ANOVA for (g to j) or two-way ANOVA for (B to D) (*p < 0.05; ** p < 0.01; *** p < 0.001; **** p < 0.0001). ns, no significance.

To further support a role for autophagy in the regulation of MAVS, we monitored MAVS levels upon depletion of autophagy genes ATG7 or ATG9A, in the presence or absence of BafilomycinA1, the autophagosome fusion blocker which prevents autophagy-dependent degradation of cargo, and thus allows accumulation of cargo ^68^. We found that the basal level of MAVS was increased upon treatment with Bafilomycin A1 (Fig. 4e). Furthermore, the increased levels of MAVS observed upon autophagy depletion remained high with BafilomycinA1 treatment (Fig. 4e). These data suggest that MAVS is negatively regulated by autophagy.

MAVS is mitochondria-associated, and activation occurs on the mitochondrial membrane impacting mitochondrial morphology ^6,65,66^. Furthermore, ATG9A is associated with autophagy at mitochondria further suggesting a direct role ^69^. Therefore, we used confocal microscopy to monitor the impact of autophagy genes on MAVS and mitochondria. As expected, basally we observed high co-localization between MAVS and mitochondria as measured by TOMM20 (Extended Data Fig. 5a, and 5b). Upon depletion of autophagy genes ATG7 or ATG9A, we found significant increases in the intensity of MAVS as well as increases in the mitochondrial branch length and branch network, features associated with signaling (Fig. 4f to i, and Extended Data Fig. 5c). Altogether demonstrating that autophagy negatively regulates MAVS at mitochondria.

### SARS-CoV-2 ORF9b overcomes RLR pathway signaling

SARS-CoV-2 antagonizes early detection and thus IRF3 translocation is not observed in infected cells (Fig. 3o and p). Ectopic expression studies in non-respiratory cells have shown that a number of coronavirus proteins (NSP5, N, M and ORF9b) can bind to MAVS and inhibit IFN production ^26,40,44,47,50,70,71^. Given our observation that MAVS is specifically regulated by autophagy, we set out to determine which of these viral proteins co-localize with MAVS during bona fide SARS-CoV-2 infection of Calu-3 cells. We found that only ORF9b clearly co-localizes with MAVS during infection (Fig. 4j, k and Extended Data Fig. 5d). These results suggest that ORF9b interacts with MAVS, and that this blocks IFN signaling in SARS-CoV-2 infected cells. However, upon loss of autophagy increased MAVS levels overcome ORF9b-mediated attenuation allowing IRF3 activation in infected cells.

### SARS-CoV-2 variants express high levels of ORF9b

Since its introduction in 2019 SARS-CoV-2 has continued to adapt to humans. While Spike adaptations to overcome pre-existing antibodies has been clearly established, the trajectory of other changes is less well understood ^54^. Some studies have suggested that particular variants, for example Alpha, has increased innate immune evasion due to increased expression of accessory genes including ORF9b ^44^. Whether additional variants including Omicron lineages that have swept globally have increased accessory gene expression or antagonism of MAVS is unknown. Therefore, we determined the levels of coronavirus proteins across a panel of variants. We probed the non-structural protein NSP1 since it is the first protein produced from the polyprotein ORF1a. We also measured the levels of the nucleocapsid protein N, as well as the accessory proteins ORF3a, ORF6, and ORF9b. We found that the ancestral variant (WA1), the Alpha variant as well as the Omicron variants BA.1 and BA.5 expressed similar levels of NSP1, N and the accessory proteins ORF6 (Fig. 5a). While BA.5 expressed reduced levels of ORF3a. Strikingly, all variants including the Omicron variants BA.1 and BA.5 expressed more ORF9b than the ancestral strain (Fig. 5a).

**Fig. 5.**
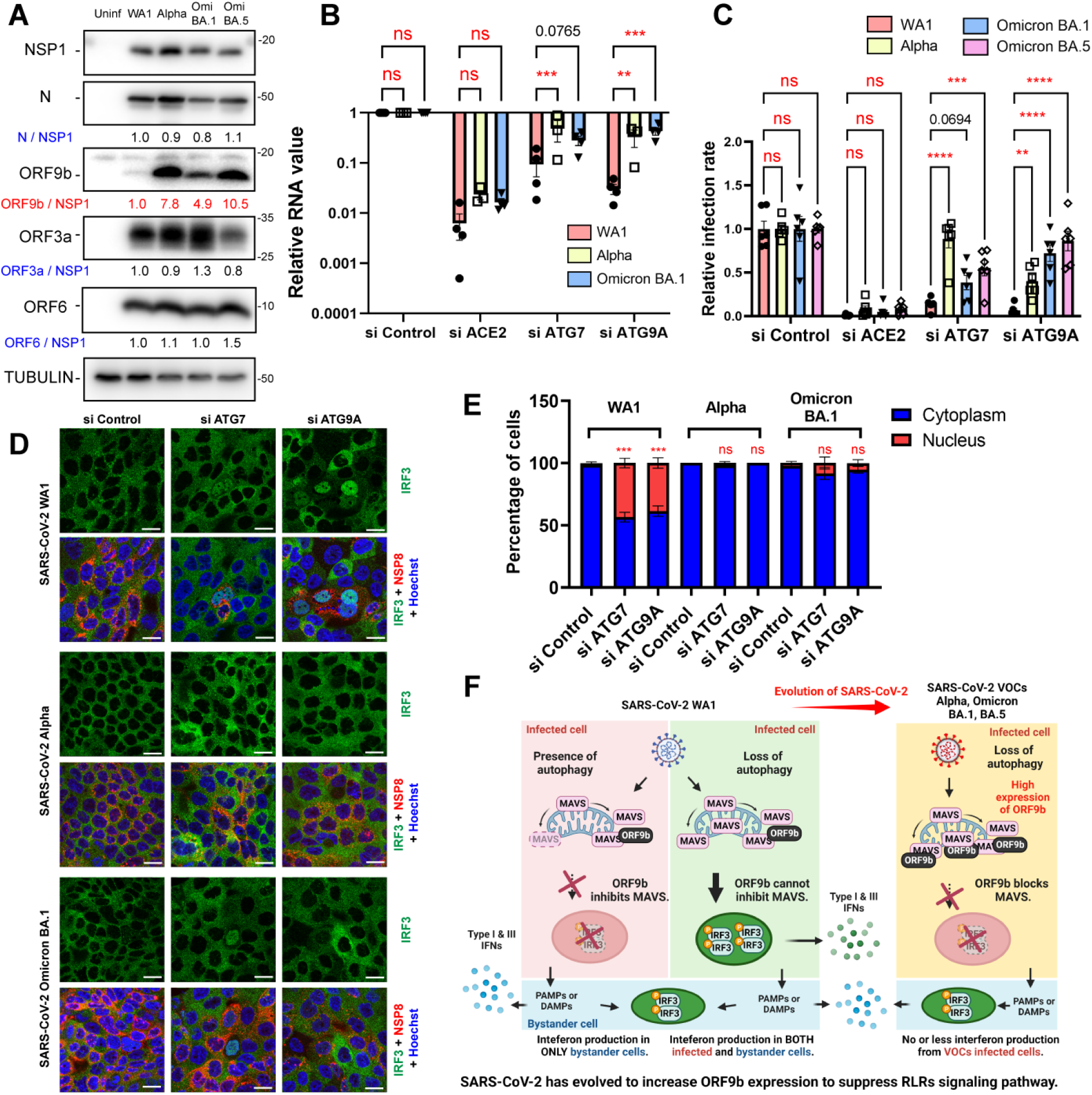
SARS-CoV-2 evolved to overcome RLR pathway via increased expression of ORF9b. (**a**) Calu-3 cells were infected with SARS-CoV-2 WA1, Alpha, and Omicron variants (BA.1, and BA.5). 2 days post-infection, total protein analyzed by immunoblot with the indicated antibodies. The intensities of viral protein normalized to NSP1 are shown. Representative blots shown for n=3. **(b)** Calu-3 cells were transfected with the indicated siRNAs and at 48 hr infected with SARS-CoV-2 WA1, Alpha and Omicron variants (MOI of 0.05). At 2 dpi, the cells were subject to RT-qPCR analysis of viral RNA (N mRNA) compared to siControl. **(c)** Calu-3 cells were transfected with the indicated siRNAs in 96 well plate, and infected with SARS-CoV-2 WA1, Alpha and Omicron variants (MOI of 0.3) at 48 hr post-transfection. After 2 days, the cells were processed for automated microscopy and automated image analysis quantified total cell numbers (nuclei+) and the percent infection (N+/nuclei+). **(d and e)** Calu-3 cells were transfected with the indicated siRNAs, 2 days post transfection, infected with SARS-CoV-2 WA1, Alpha, and Omicron variants for 2 days. The cells were fixed and processed for confocal microscopy (anti-IRF3, green; anti-NSP8, red; nuclei, Hoechst 33342, blue). Infection and nuclear IRF3 translocation were quantified. The scale bar represents 20 μm. For all graphs, data shown are means ± SEMs from biological replicates. Significance was calculated using one-way ANOVA for (f) or two-way ANOVA for (a, c and d) (*p < 0.05; ** p < 0.01; *** p < 0.001; **** p < 0.0001). ns, no significance. (**f**) Schematic describing the role of autophagy in SARS-CoV-2 infection and innate immune signaling in infected and bystander cells. SARS-CoV-2 viral RNA can be sensed by RLRs which stimulates MAVS activation. SARS-CoV-2 ORF9b evades this activation, preventing IRF3 activation in infected cells. In contrast, in uninfected bystander cells IRF3 translocates inducing IFNs and ISGs. Autophagy negatively regulates MAVS, and thus loss of autophagy increases MAVS overcoming antagonism by the ancestral strain of SARS-CoV-2. Under these conditions, IFN induction occurs both in autophagy-deficient infected cells and bystander cells blocking SARS-CoV-2 replication. However, SARS-CoV-2 variants evolved to express increased ORF9b to prevent IRF3 translocation in cells with high levels of MAVS and RLR signaling. This allows Alpha and Omicron variants to more efficiently evade innate immunity and interferon signaling.

ORF9b and N are expressed from the same subgenomic (sg)RNA, but ORF9b is expressed from a downstream start codon in a distinct reading frame. We observed increased ORF9b expression relative to N and thus we also monitored the expression of the sgRNA that encodes both proteins, and compared this to other sgRNAs. We found sgRNA encoding ORF3a was not changed across the variants, and ORF6 sgRNA was modestly reduced in BA.5. In contrast, the sgRNA encoding both N and ORF9b is significantly increased in Alpha, Omicron BA.1 and BA.5, with BA.5 expressing the most RNA and protein (Extended Data Fig. 6a).

### SARS-CoV-2 variants are resistant to autophagy-dependent control of IFN

The ancestral strain of SARS-CoV-2 expressed low levels of ORF9b that could not overcome the increased MAVS levels that accumulate upon loss of autophagy. Thus, loss of autophagy genes blocked infection by the ancestral strain. Since the more recent variants express higher levels of ORF9b, we hypothesized that these strains would be resistant to control by autophagy-dependent MAVS regulation. To test this, we depleted the autophagy genes ATG7 or ATG9A and infected with the ancestral strain, Alpha or Omicron BA.1. As a positive control, we found that all three strains were dependent on the entry receptor ACE2 (Fig. 5b). While the ancestral strain was sensitive to depletion of ATG7 or ATG9A showing significant reduction in infection, the Alpha and Omicron variants are insensitive to autophagy-gene depletion, replicating to control levels as measured by RT-qPCR (Fig. 5b). We used a microscopy-based assay to corroborate these data. Depletion of ACE2 blocked replication of all strains tested (ancestral strain, Alpha, Omicron BA.1 and Omicron BA.5) (Fig. 5c). Moreover, depletion of ATG7 or ATG9A led to a striking reduction in infection by the ancestral strain WA1 (Fig. 5c). In contrast, loss of autophagy genes had minimal impact (< 2-fold) on the more recent variants (Fig. 5c). Altogether, this suggests that Alpha and Omicron strains can overcome higher levels MAVS and suppress IFN signaling.

To test whether Alpha and Omicron could indeed suppress IRF3 activation in autophagy-depleted infected cells we performed confocal microscopy. As expected, in control-depleted cells WA1, Alpha and Omicron suppress IRF3 translocation in infected cells (Fig. 5d to e). Upon loss of autophagy genes, the ancestral WA1 strain can no longer suppress IRF3, and nuclear IRF3 is now observed in infected cells. (Fig. 5d to e). In contrast, Alpha and Omicron variants have little nuclear IRF3 translocation in autophagy-depleted infected cells (Fig. 5d, and e). Altogether, these data suggest that evolution of increased expression of ORF9b promotes antagonism of higher levels of RLR signaling during infection.

## Discussion

IFN signaling plays a fundamental role in controlling viral infections generally, and in particular against SARS-CoV-2 infection ^4,12,31,35,44,45,61,62^. Individuals with either loss-of-function mutations in genes required for IFN induction or auto-antibodies to IFNs show increased mortality ^8,9^. Since SARS-CoV-2 evades early detection in respiratory epithelial cells, we suggest that this evasion promotes the establishment of infection in the respiratory tract ^10,12,39–44,70^. Increasing early detection or inducing a more rapid IFN responses at barrier sites will likely ameliorate disease ^67^. Indeed, early treatment with IFNs or STING agonists that induce IFNs can protect from infection ^8,12,46,48^. However, while IFN is important during early phases it can be detrimental during late phases. For example, severe COVID19 patients can present with high IFN and inflammatory responses which may also contribute to disease ^1,9,72^. Therefore, the balance of IFN signaling is crucial. Here we show that autophagy regulates the basal levels of MAVS, an essential component of innate immune signaling. In autophagy-deficient cells increased MAVS leads to IFN stimulation; and this can prevent SARS-CoV-2 infection.

Autophagy plays homeostatic roles to maintain the balance of many cellular processes largely related to stress ^36^. Furthermore, autophagy can play diverse roles in viral infections, having direct roles in viral replication and indirect roles in regulating pathways that impact infection including immunity ^13,22^. Previous studies of SARS-CoV-2 infection in non-respiratory cells implicated autophagy in promoting early replication steps ^24,25,73–75^. We also found that autophagy promotes infection; however, this requirement for autophagy is not to promote replication directly but rather to restrain interferon signaling. We found that in the absence of interferon signaling autophagy genes are dispensable for SARS-CoV-2 infection (Fig. 2J and Extended Data Fig. 3e to 3g). This led us to explore the relationship between autophagy genes and IFN signaling in the context of SARS-CoV-2 infection.

Homeostasis of innate immune signaling is essential as dysregulation of IFNs resulting in high basal levels can cause diverse human diseases from inflammatory and autoimmune to neurodegeneration ^62,76,77^. Therefore, it is not surprising that many of signaling molecules involved in innate immunity including RIG-I, MDA5, STING, MAVS, TBK1, and IRF3, can be regulated by autophagy under some conditions ^33–37,61,78^ ^26^. We found that autophagy genes involved in each step of the autophagy pathway impacted SARS-CoV-2 infection. Moreover, we found that autophagy genes and factors specifically involved in mitophagy also impact infection and innate immune signaling. For example, while diverse cargo receptors such as SQSTM1 (p62), CALCOCO2 (NDP52), OPTN, and NBR1 ^34,35,37,79–81^ can play roles in mitophagy, we found that only OPTN had striking roles in SARS-CoV-2 replication impacting basal IFN signaling. Furthermore, recent studies suggested that ATG9A along with OPTN control mitophagy ^19–21,82^. We found that ATG9A also impacted SARS-CoV-2 replication. While previous studies linked ATG9A to the regulation of STING, we found that the activity of autophagy genes is STING-independent (Fig. 2k, and Extended Data Fig. 3i to 3k) ^33^. A recent study found that ATG9A and OPTN impacts components of the PRR pathway upstream of IFN, this was not through MAVS or dependent on canonical autophagy genes ^83^. In this study, we found that genes involved in all steps of mitophagy, from cargo recognition to late steps in auto-lysosomal fusion (STX17, VAMP8, VSP39, and VPS41) control IFN and SARS-CoV-2 replication. Altogether, these data suggest that the full cycle of autophagy impacts SARS-CoV-2 replication through regulation of MAVS and IFN production.

Mitochondria play a major role in innate immunity and the RLR pathway since MAVS is localized and activated at mitochondria ^62,82^. Furthermore, studies have shown that increased MAVS levels can drive increased IFN activity even in the absence of viral infection ^6,84,85^. Thus, MAVS levels must be tightly regulated. Furthermore, we found that unlike many components of this signaling pathway MAVS is not an ISG but rather controlled by autophagy (Fig. 4a to e). While MAVS-dependent IFN induction is important for early control of SARS-CoV-2 ^53,71^, SARS-CoV-2 can evade early activation ^4,12,45^. Many studies have explored the role of SARS-CoV-2 proteins as innate immune antagonists using ectopic expression finding that many proteins can block signaling ^40–44,47–50,70,71^. Furthermore, interactome studies have also found that many viral proteins can interact with MAVS or mitochondrial proteins ^86–88^. For example, studies have shown that MAVS activity can be regulated by N, M, NSP5, ORF9b, and ORF10 ^41,44,50,70,71^. This led us to determine which of these proteins, when expressed during bona fide infection in respiratory cells, co-localized with MAVS. We found that only ORF9b has high co-localization pattern with MAVS (Fig. 4j, k, and Extended Data Fig. 5d) and thus we further explored this biology.

Viral antagonists, such as ORF9b, function to block innate immune signaling in the infected cell. And if successful, leads to innate activation of IFNs and ISGs only in uninfected bystander cells ^12,45^. This leads to delayed responses which can promote spread of the virus before innate immune control. We found that basal autophagy maintains low levels of MAVS which can be suppressed by ORF9b in infected cells to prevent IRF3 translocation and IFN production in infected cells. Bystander cells then recognize viral products released from infected cells. However, upon loss of autophagy genes MAVS levels are increased, overcoming ORF9b, leading to IRF3 activation and IFN production in infected cells to block SARS-CoV-2 replication. Thus, under these increased MAVS conditions both infected and uninfected cells are responding.

This led us to explore SARS-CoV-2 variants as SARS-CoV-2 is continuing to adapt successfully infecting humans in the face of pre-existing immunity ^54,57,89–91^. In addition to evasion of antibodies, recent studies have suggested that additional adaptations have led to decreased sensitivity to innate immune control ^44,92^. In particular a recent study showed that the Alpha variant contains a point mutation generating partial transcriptional regulatory sequence (TRS) for sgRNA of ORF9b, so it leads to increased ORF9b expression ^44^. While Omicron variants including BA.1 and BA.5 do not harbor this mutation, we found that these variants express even higher levels of ORF9b (Fig. 5a and Extended Data Fig. 6a). It remains unclear how Omicron variants induce higher levels of the sgRNA that expresses ORF9b. Nevertheless, these data suggest that there is a selective pressure to increase ORF9b to block innate immune activation. Altogether, these studies provide mechanistic insight into how autophagy regulates IFN signaling to control viral infections and the selective pressure this is placing on SARS-CoV-2 to evolve to suppress interferon signaling to continue to gain additional footholds in human populations.

## Acknowledgments

We thank members of the Cherry, Lynch and UPENN High-Throughput Screening Core, for technical advice and discussion; Emily Lee and Marc Ferrer, NCATS, and Mattek for the bronchial ALI cultures; Samuel Constant and Epithelix for the nasal ALI cultures; and Wenli Yang and the iPSC Core at UPENN for the iAT2 cells. We thank Don Pijak and EHRS for BSL3 maintenance.

This work was supported by grants from the National Institutes of Health to S. Cherry (1-R01-AI-150246, 1-R01-AI-152362 1-R01-AI-140539), Mark Foundation (19-011MMIA) as well as funding from the Penn Center for Precision Medicine, Mercatus and the Bill and Melinda Gates Foundation (INV-018479). S. Cherry was a recipient of the Burroughs Wellcome Investigators in the Pathogenesis of Infectious Disease Award, the Deans Innovation Fund as well as Linda and Laddy Montague.

## Author contributions

Conceptualization: J.S.L., S.C.

Methodology: J.S.L., M.D., M.L., M.F., T.G., D.C.S., S.C.

Validation: J.S.L, J.M., K.A., J.H., K.W., D.S.C., S.C.

Formal Analysis: J.S.L, J.M., K.A., J.H., K.W., D.S.C., S.C.

Investigation: J.S.L., M.D., J.M., K.A., M.F., K.W., Z.E., D.S.C.

Resources: D.S.C., S.C.

Data Curation: J.S.L, M.F., D.S.C., S.C

Writing – original draft: J.S.L., S.C.

Writing – review & editing: S.C. M.D., T.G, D.C.S, K.A.

Visualization: J.S.L., T.G.

Supervision: S.C.

Project administration: S.C.

Funding acquisition: S.C.

## Methods

### Cell culture

Human adenocarcinoma lung epithelial cells, Calu-3 (ATCC, HTB-55), were cultured in MEM (GIBCO) supplemented with 10 % (v / v) fetal bovine serum (FBS), 1 % (v / v) penicillin / streptomycin, 1 % (v / v) Glutamax, and 1 % (v / v) non-essential amino acid, at 37 ℃ and 5 % CO_2_ incubator ^94^. Rat tail collagen coated plates (Corning) and pre-coated cover slips (Electron Microscopy Sciences) were used for all Calu-3 experiments. Monkey kidney cells, VERO E6 (ATTC CRL-1586) and VERO-TMPRSS2 ^59,94^, were culture in DMEM (GIBCO) supplemented with 10 % (v / v) fetal bovine serum (FBS), 1 % (v / v) penicillin / streptomycin, and 1 % (v / v) Glutamax. Day 20 air liquid interface (ALI) EpiAirway tissues were provided from MatTek Corporation (AIR-100). Tissues were fed twice in a week until use. Air liquid interface pooled nasal epithelial cultures from Epithelix were fed twice in a week until use. For ALI infections, the apical surface was washed with Opti-MEM, and cells placed into fresh media with drugs added to the basolateral surface. Cells were infected apically with SARS-CoV-2 (MOI of 0.2) for one hour and subsequently the virus inoculum was removed as described (both references). iAT2 cells were differentiated from SPC2 iPSC line, clone SPC2-ST-B2 (Boston University) and maintained as alveolospheres. The alveolospheres were dissociated into single cells and plated on 3% Matrigel coated plates in CK+DCI media supplemented with 2 μM TZV for 2 days. Culture medium was changed to CK+DCI media 3 days post-plating prior to SARS-CoV-2 infection as described (24, 55).

### Viruses

SARS-CoV-2 was obtained from BEI (WA-1 strain, Alpha, Omicron BA.1 and BA.5) ^12,94^. SARS-CoV-2 virus stocks were prepared by infecting VERO-TMPRSS2 cells with media containing 2 % FBS supplemented with HEPES. After 3 days of virus infection, the cells were freeze-thawed and media was collected and centrifuged 3,000 rpm for 20 min. The supernatant containing virus was aliquoted and stored at −80 ℃ (P0). The seed stock (P0) was sequence verified, amplified in VERO E6 cells (P1), and used for all experiments. The titer of virus stock was determined by 50% tissue culture infective doses (TCID_50_) using the Reed-Muench method in VERO E6 cells as described previously ^59^. All work with SARS-CoV-2 was performed in a biosafety level 3 (BSL3) laboratory and approved by the Institutional Biosafety Committee and Environmental Health and Safety. Sendai virus was obtained from Charles River (Cantell strain). Vesicular stomatitis virus (VSV-eGFP) is a kind gift from J. Rose. The virus stock was grown in BHK cells as described ^93^.

### RNA preparation and RT-qPCR

Total RNA was purified using Trizol (Invitrogen) followed by RNA Clean and Concentrator-25 kit (Zymo Research). Random hexamers, dNTP and Moloney murine leukemia virus (M-MLV) reverse transcriptase (Invitrogen) were used to synthesize cDNA. Gene specific primers and SYBR green master mix (Applied Biosystems) were used to amplify target genes and 18S rRNA primers were used to amplify endogenous control by using the QuantStudio 6 Flex RT-PCR system (Applied Biosystems). Relative quantities of viral and cellular RNA were calculated using the standard curve method ^12,59^.

### Immunoblot

Calu-3 cells were washed with PBS and lysed by radioimmunoprecipitation assay (RIPA) buffer [50 mM tris-HCl, (pH 8.0), 150 mM NaCl, 0.5% sodium deoxycholate, 0.1% SDS, 1% NP-40, 1 mM PMSF, protease inhibitor cocktail and phosphatase inhibitor cocktail] ^12^. Upon centrifugation at 13,000 rpm for 15 min at 4 ℃, clarified cell lysates were incubated with 5X sample buffer at 95 ℃ for 10 min. Protein samples were resolved by sodium dodecyl sulfate-polyacrylamide gel electrophoresis (SDS-PAGE) followed by transfer to PVDF membrane (Millipore). The membrane was blocked with 5 % skim milk in tris-buffered saline with 0.1% Tween20 (TBS-T) for 30 min at room temperature and incubated with primary antibody overnight at 4 ℃. Membranes were washed 3 times with TBS-T, and then incubated with horseradish peroxidase (HRP)–conjugated secondary antibody for 2 hr at room temperature. The membrane was washed 3 times with TBS-T and developed with West Femto Substrate (Thermofisher) and ECL Western blotting reagents (Amersham). Western blotting signals were visualized by using Amersham Imager 680 (Amersham).

### Immunofluorescence assay

Calu-3 cells were plated on collagen coated cover slips and treated as indicated ^12^. The cells were fixed with 4 % formaldehyde for 15 min at room temperature and rinsed 3 times with PBS. The cells were permeabilized and blocked by 2 % BSA in PBS with 0.1 % Triton X 100 (PBS-T) for 30 min at room temperature. The coverslip was incubated with primary antibody for overnight at 4 ℃. The coverslip was rinsed 3 times with PBS-T and incubated with Alexa Flour conjugated secondary antibodies (Sigma-Aldrich) and Hoechst 33342 (Sigma-Aldrich) for 1 hr at room temperature. After 3 washes, the coverslip was mounted on slide glass with VECTASHIELD Antifade Mounting Medium (Vector Laboratories) and sealed with nail polish. The samples were visualized by Leica DM5500Q confocal microscope at 20X or 63X. Nuclear translocation of IRF3 was analyzed by the NuclearHTtranslocation module in MetaXpress 5.3.3 software.

### Viral titration and automated microscopy

Automated microscopy experiments were performed as described previously ^59^. Briefly, Calu-3 cells were plated on collagen coated 96 well plates and treated as indicated. Next, the cells were fixed with 4 % formaldehyde for 15 min at room temperature and washed 3 times with PBS. The cells were permeabilized and blocked by 2 % BSA in PBS-T for 1 hr at room temperature. The plate was incubated with primary antibody for overnight at 4 ℃, washed 3 times with PBS-T, and incubated with Alexa Flour conjugated secondary antibodies and Hoechst 33342 for 1 hr at room temperature. After 3 washes, the plate was sealed and imaged using an automated microscope (ImageXpress Micro, Molecular Devices). Cells were imaged with a 10X objective, and 4 sites per well were captured. The total cell numbers and the infected cell numbers were measured using the cell scoring software (MetaXpress 5.3.3), and the percentage of infected cells was calculated. Titer of virus stock was determined by 50% tissue culture infective doses (TCID_50_) of Reed-Muench method in VERO-TMPRSS2 or Calu-3 cells.

### siRNA transfection

The gene of interest expression was depleted by transfection with siRNAs using Lipofectamine RNAiMax reagent (Thermo Scientific) according to manufacturer’s protocol. Calu-3 cells were plated on 6 well plate (5 × 10^5^), 96 well plate (4.5 × 10^4^) or coverslips (2 × 10^5^) containing an siRNA: transfection reagent mixture. Two siRNAs for each selected host target were co-transfected and the media were changed after overnight incubation. 2 days post transfection, the cells were inoculated with SARS-CoV-2 (MOI of 0.2). After 48 hr infection, the cells were collected with either TriZol to analyze RNA levels, or with RIPA buffer to be used for western blotting, or fixed with 4 % formaldehyde for measuring the percentage of infection via automated microscopy.

### SARS-CoV-2 virion particle production measurement from the knock down cells

Titration experiments were performed as described previously ^59^. Calu-3 cells were transfected with target siRNAs on 6 well as described above and inoculated with SARS-CoV-2 after 2 days post transfection. The media was changed at 4 hr post infection. Two days later, 1 ml of supernatants were collected and centrifuged at 5000 rpm for 30 min at 4 ℃. The supernatants were collected and kept on −80 ℃. For measuring virus titer, TCID_50_ of Reed-Muench method was used with VERO-TMPRSS2 cell as described above (*24*).

### Dose response analysis for VSV

Dose response experiments were performed as described previously (*24*). Calu-3 cells (7.5 × 10^3^) were plated in collagen-coated 384 well plates (Corning BioCoat). Next day, cells were treated with 0.2 % DMSO, serial diluted VSP34 IN1 in 0.2% DMSO or IFNβ. After 2 hr, the cells were inoculated with VSV-GFP (MOI of 1.8) (*56*). The cells were fixed at 24 hpi in 4 % formaldehyde and then washed three times with PBS. The cells were permeabilized and blocked by incubating with 2 % BSA in PBS-T for 1 hr at room temperature. The plate was incubated with Hoechst 33342 for 1 hr at room temperature. After 3-time washes, the plate was sealed and imaged using an automated microscope (ImageXpress Micro, Molecular Devices). Cells were imaged with a 10X objective, and 4 sites per well were captured. The total cell numbers and the infected cell numbers were measured using the cell scoring software (MetaXpress 5.3.3), and the percentage of infected cells was calculated.

### Co-localization and MAVS morphology analysis

Calu-3 cells were plated on collagen coated cover slips and transfected with indicated siRNAs. Two days post transfection, the cells were inoculated with SARS-CoV-2 and fixed at 24 hpi. Fixation was performed with 4% formaldehyde for 10 min and washed 3 times with PBS. Viral proteins were stained with specific antibodies with MAVS antibody (sc-166583). To analyze co-localization coefficient, coloc2 plug-in was used in Image J software. For MAVS morphology analysis, MAVS protein was stained with MAVS antibody (sc-166583) and TOMM20 antibody (ab56783) was used for staining mitochondria. MAVS signal intensity was measured by ImageJ software. MAVS and mitochondrial morphologies were analyzed by Mitochondrial Network Analysis (MiNA) toolset in ImageJ ^96^.

### Immunoblot analysis of SARS-CoV-2 variants of concern

Calu-3 cells were infected with SARS-CoV-2 variants of concern (WA1 (MOI 0.1), Alpha (MOI 0.1), Omicron BA.1 (MOI 0.3), and BA.5 (MOI 0.1). After 2 days post infection, the cells were lysed and processed for immunoblot. The intensity of viral proteins was normalized to NSP1 protein which is coded in 5’ end of ORF1ab. Equal levels of replication are observed as measured by NSP1 levels.

### Subgenomic RNA level of SARS-CoV-2 variants

Calu-3 cells were infected with SARS-CoV-2 variants of concern (WA1 (MOI 0.1), Alpha (MOI 0.1), Omicron BA.1 (MOI 0.1), and BA.5 (MOI 0.1). for 2 days. Total RNA was collected and processed for measuring qPCR. To detect specific subgenomic RNA (sgRNA), the same forward TRS primer (sgRNA-F) and specific reverse primer for each sgRNA region were used (sgRNA-ORF3a-R, sgRNA-ORF6-R, and sgRNA-N-R) ^97^. Each sgRNA level was normalized by genomic RNA level which was measured by sgRNA-F and 5’UTR-R primers.

### Quantification and statistical analysis

Statistical analyses were performed by using Prism (GraphPad Software, V9.4.1). Statistical significances were investigated by unpaired t test, ordinary one-way ANOVA or two-way ANOVA with multiple comparisons for 95% CI value. Adjusted p values are described by asterisks in figures: (*) for p < 0.05, (**) for p < 0.01, (***) for p < 0.001, and (****) for p < 0.0001.

## Supplemental Information

Supplemental table 1 to 3

## Extended Data

**Extended Data Fig. 1.**
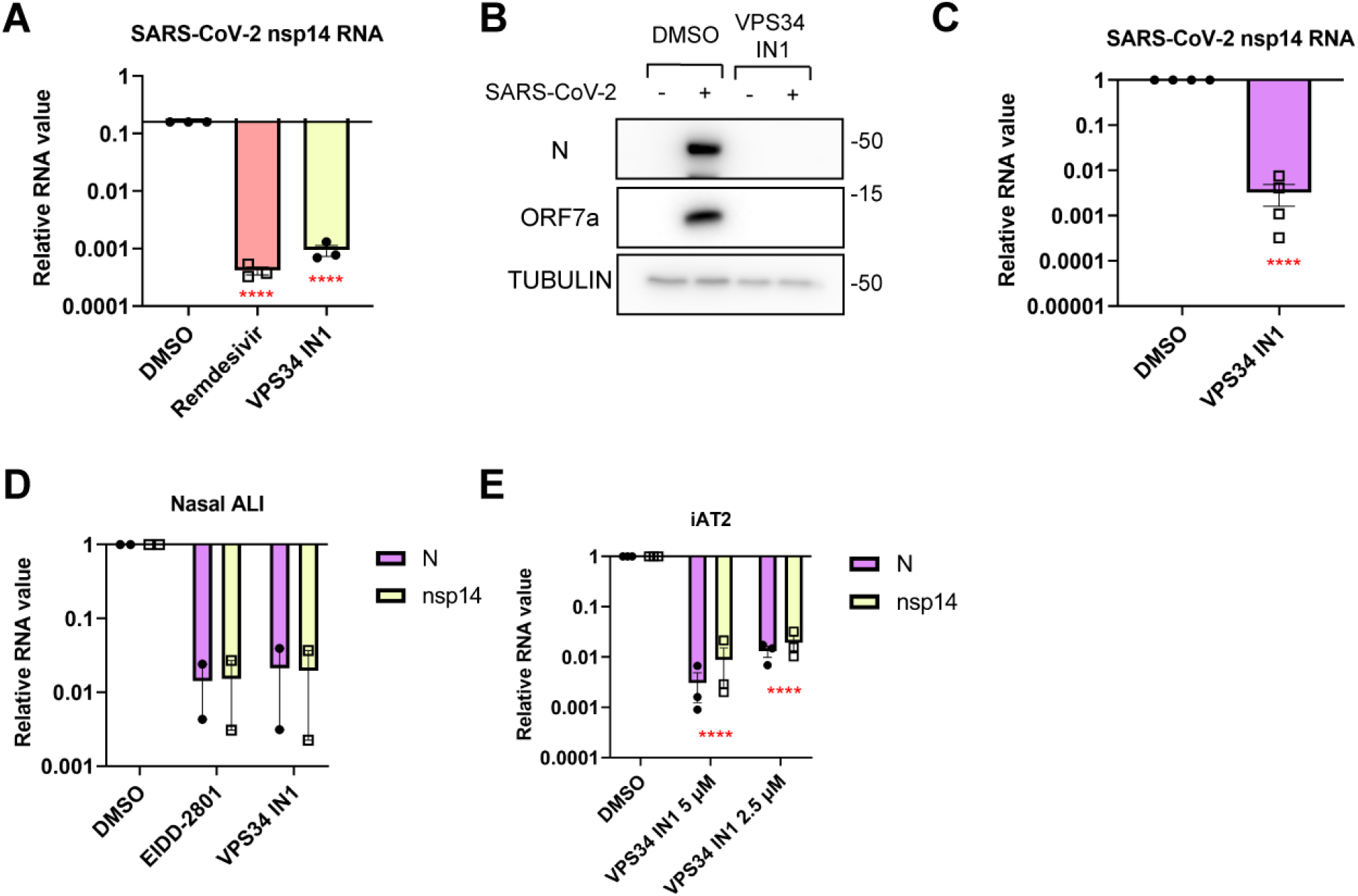
VPS34 IN1 controls SARS-CoV-2 infection in diverse respiratory models. (**a** and **b**) Calu-3 cells were pretreated with vehicle (DMSO) or the indicated drug at 10 μM, followed by infection with SARS-CoV-2 (MOI of 0.2). At 48 hr post infection (hpi), total RNA or total protein was isolated for (a) RT-qPCR against viral RNA (nsp14) or (b) immunoblot with the indicated antibodies. Representative blots shown for n=3. (**c**) Human bronchial air-liquid interface (ALI) cells were pretreated with vehicle or 3 μM VPS34 IN1 and then infected with SARS-CoV-2 (MOI of 0.2). Viral RNA levels were measured at 72 hpi (nsp14). (**d**) Primary human nasal epithelial cells in air liquid interface culture (ALI) were pretreated with vehicle or the indicated drugs at 3 μM followed by infection with SARS-CoV-2 (MOI of 0.2). Viral RNA levels were analyzed at 72 hr post-infection (hpi) using RT-qPCR for the indicated viral targets (N and nsp14). (**e**) human iPSC derived iAT2 cells were treated with vehicle or the indicated concentrations of VPS34 IN1 and infected with SARS-CoV-2 (MOI of 0.2) for 72 hr. Total RNA was collected to measure viral RNA levels by RT-qPCR for the indicated viral target. For all graphs, data shown are means ± SEMs from indicated replicates. Significance was calculated using unpaired t-test for (b) one-way ANOVA for (a), or two-way ANOVA for (e) (**** p < 0.0001).

**Extended Data Fig. 2.**
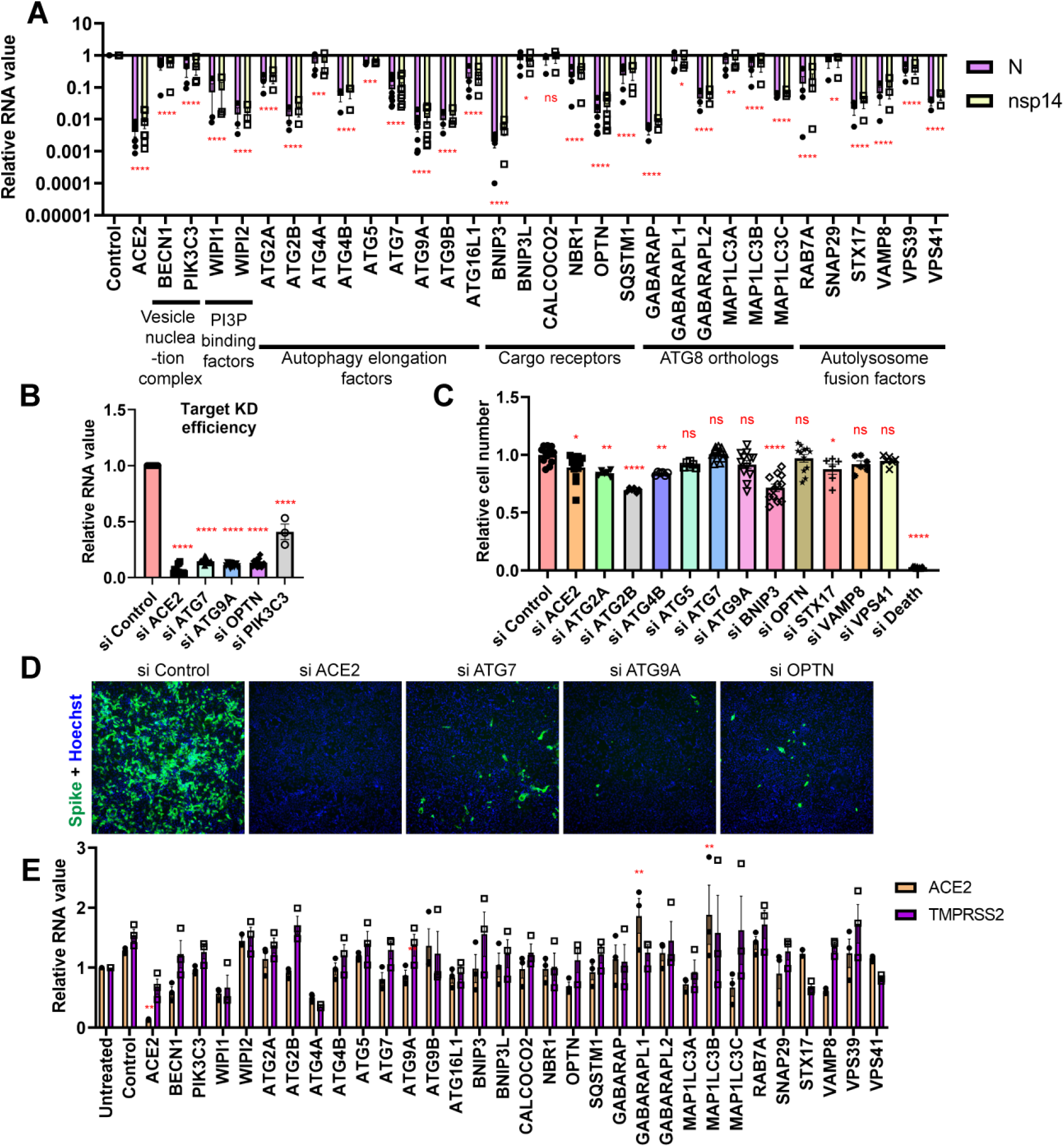
Autophagy genes promote SARS-CoV-2 infection. (**a** and **b**) Calu-3 cells were transfected with the indicated siRNAs and 48h later infected with SARS-CoV-2 (MOI of 0.2). At 48 hpi, viral RNA levels were quantified by RT-qPCR against the indicated viral targets or cellular targets shown relative to siControl. (**c** and **d**) Calu-3 cells were transfected with the indicated siRNAs, and 2 days later, infected with SARS-CoV-2 (MOI of 0.5). After 48 hr, the cells were processed for automated microscopy and automated image analysis (Spike, green) and total cells (Hoechst 33342, blue). We quatified total cell number and the percent infection. Representative images shown. (**e**) mRNA level of ACE2 and TMPRSS2 was determine by qPCR in the indicated siRNA transfected samples from Fig. S2A. (**) represents p < 0.01 in ACE2 mRNA. For all graphs, data shown are means ± SEMs from biological replicates. Significance was calculated using one-way ANOVA for (b, c, and e) or two-way ANOVA for (a) (*p < 0.05; ** p < 0.01; *** p < 0.001; **** p < 0.0001). ns, no significance.

**Extended Data Fig. 3.**
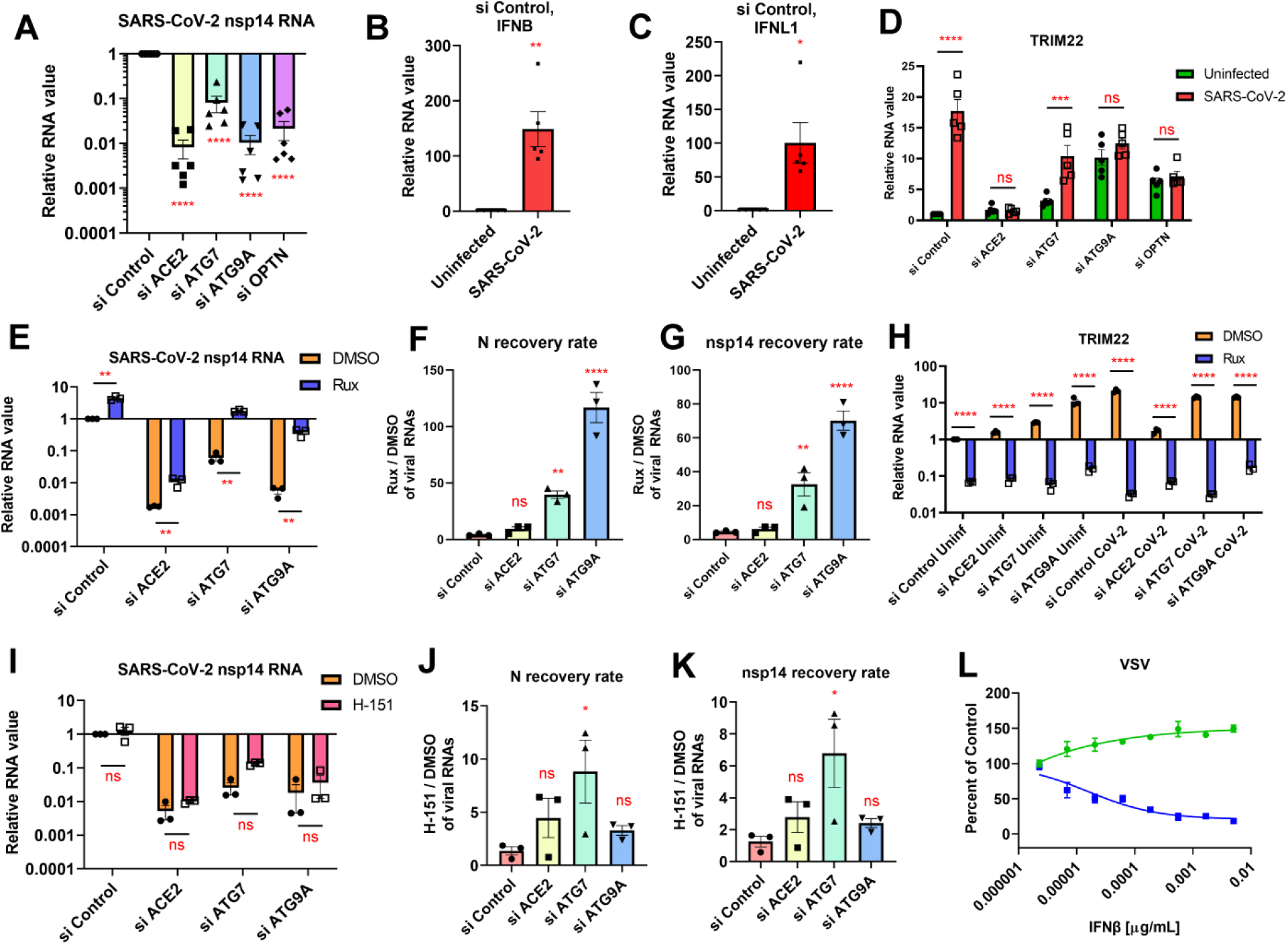
Autophagy controls SARS-CoV-2 infection through interferon signaling. (**a** to **d**) Calu-3 cells were transfected with the indicated siRNAs, and 48h later either uninfected or infected with SARS-CoV-2 (MOI of 0.2). At 48 hpi, total RNA was collected and subjected to RT-qPCR against the indicated targets. (**e** to **k**) Calu-3 cells were transfected with the indicated siRNAs, after 2 days, cells were treated with vehicle or 10 μM Ruxolitinib (JAK inhibitor) (e to h) or 10 μM H-151 (STING inhibitor) (i to k) and infected with SARS-CoV-2 (MOI of 0.2) for 48 hr. Viral genomic RNA level was analyzed by RT-qPCR against the indicated viral target (N or nsp14) and normalized to vehicle-treated siControl transfected infected sample. ISG TRIM22, level was also measured and data normalized to vehicle-treated control siRNA transfected uninfected cells (**l**) Calu-3 cells were pre-treated with vehicle or IFNβ at the indicated concentration and infected with VSV-GFP (MOI of 1.8). At 24 hpi, the cells were fixed with 4 % formaldehyde and the percentage of infection was measured by automated microscopy with cell number (green) and percent infection (blue) shown as percent of vehicle control. For all graphs, data shown are means ± SEMs from biological replicates. Significance was calculated using unpaired t-test for (b and c), one-way ANOVA for (a, d, e, h, and i) or two-way ANOVA (b, c, f, and g) (*p < 0.05; ** p < 0.01; *** p < 0.001; **** p < 0.0001). ns, no significance.

**Extended Data Fig. 4.**
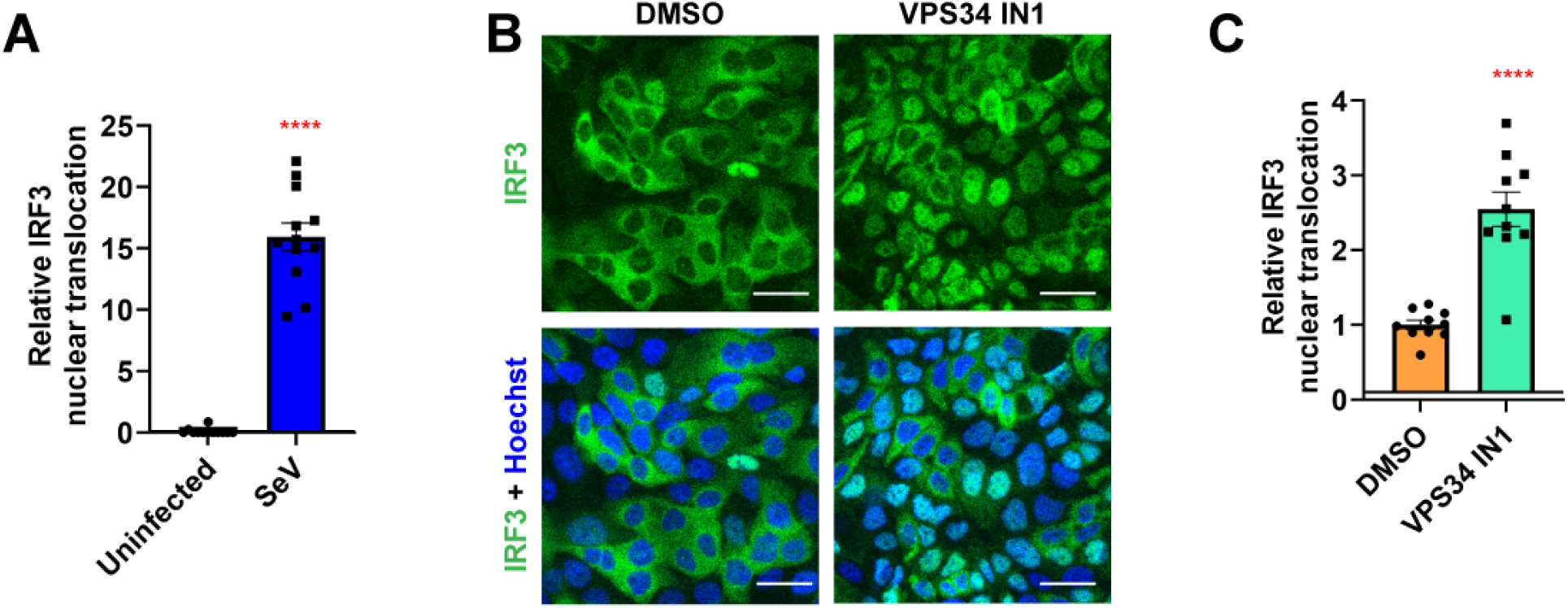
VPS34 inhibitor induces IRF3 translocation during SeV infection. (**a**) Calu-3 cells were inoculated with Sendai virus (SeV, 100 HAU / ml) for 6 hr and the cells were processed and analyzed for the relative IRF3 nuclear translocation compared to uninfected cells. Two independent experiments were performed, each set has at least 5 images and each dot represents a single analyzed image. (**b** and **c**) Calu-3 cells were pretreated with vehicle or 2 μM VPS34 IN1 and inoculated with Sendai virus (SeV, 100 HAU) for 6 hr. The cells were fixed and processed for IRF3 immunostaining (anti-IRF3, green; nuclei, Hoechst 33342, blue) and quantified for nuclear IRF3 translocation normalized to vehicle control. The white scale bar represents 40 μm. Three independent experiments were performed, each set has at least 3 images and each dot represents a single analyzed image. Data shown are means ± SEMs from described replicates. Significance was calculated using unpaired t-test (**** p < 0.0001).

**Extended Data Fig. 5.**
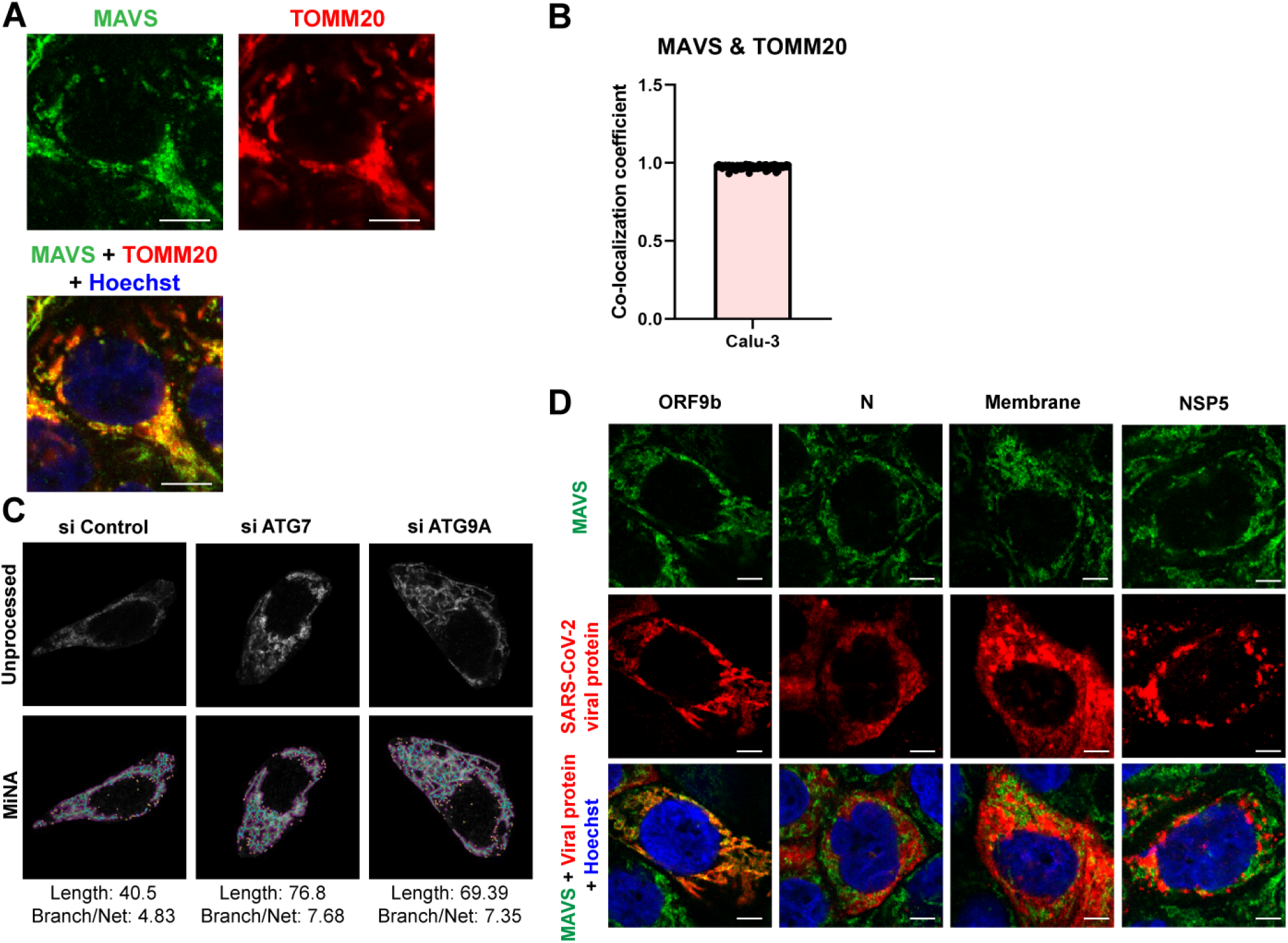
Mitochondrial MAVS co-localizes with ORF9b in SARS-CoV-2 infected cells. (**a** and **b**) Calu-3 cells were fixed and processed for confocal microscopy. Representative images shown (anti-MAVS, green; mitochondria, anti-TOMM20, red; nuclei, Hoechst 33342, blue). Co-localization coefficient was determined for each cell via using Coloc2 tool in Image J software. Three independent experiments were performed and more than 67 cells were analyzed. The scale bar represents 10 μm. Data shown are means ± SEMs from indicated replicates. (**c**) Calu-3 cells were transfected with the indicated siRNAs for 3 days and processed for confocal microscopy. MAVS localization pattern of each cell was analyzed by MiNA toolset of Image J software for accumulated length, and number of branches per network, representative mask shown. (**d**) Calu-3 cells were infected with SARS-CoV-2 (0.2 MOI) for 24h and processed for microscopy (anti-MAVS, green; anti-ORF9b, anti-nucleocapsid, anti-membrane, anti-NSP5, red; nuclei, Hoechst 33342, blue). The scale bar represents 5 μm.

**Extended Data Fig. 6.**
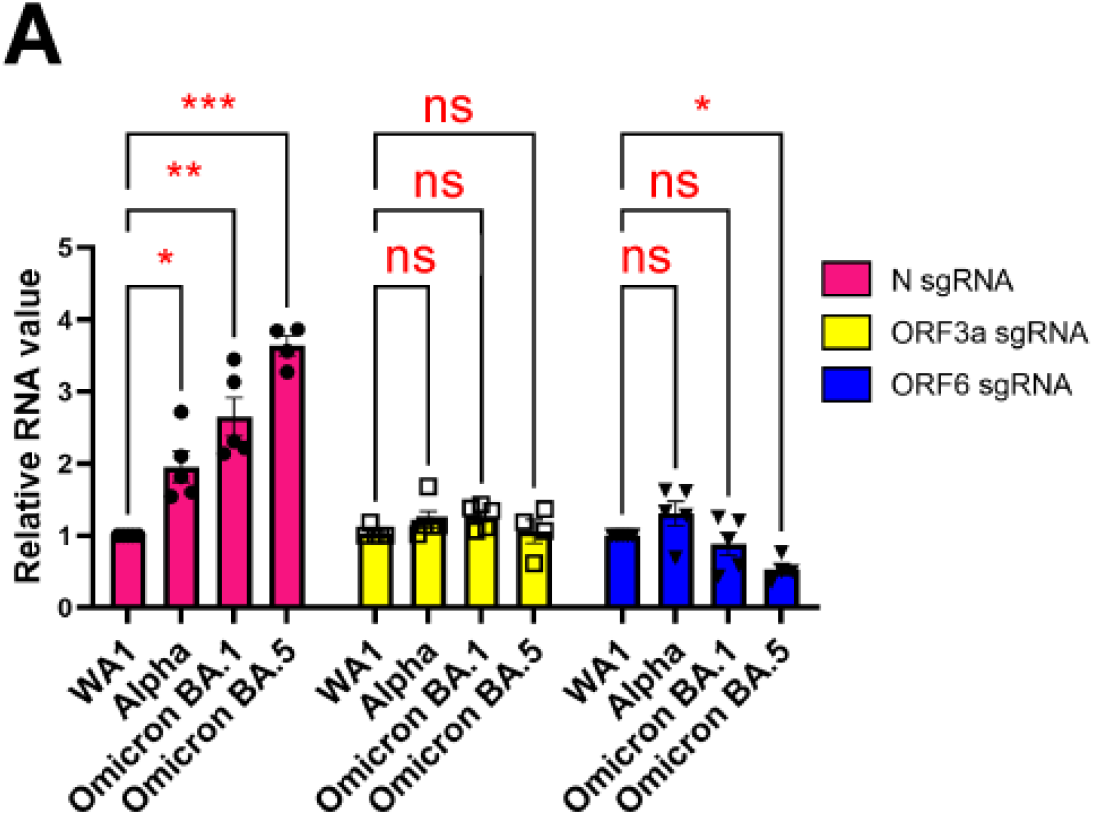
Expression level of SARS-CoV-2 subgenomic RNAs across variants of concern. (a) Calu-3 cells were infected with SARS-CoV-2 WA1, Alpha, and Omicron variants (BA.1, and BA.5). 2 days post-infection, total RNA was subjected to RT-qPCR analysis of viral subgeomic mRNA compared to WA1. Each sgRNA level was normalized by 5’UTR genomic RNA levels. Significance was calculated using two-way ANOVA (*p < 0.05; ** p < 0.01; *** p < 0.001). ns, no significance.

