## Supplemental Tables for "Evolutionary arms race between SARS-CoV-2 and interferon signaling via dynamic interaction with autophagy"

### Supplemental information

#### Supplemental table 1. Reagents used in this study.

| **Reagents** | **Source** | **Identifier** |
|---|---|---|
| Antibodies | | |
| Anti-SARS-CoV-2 nucleocapsid | GeneTex | GTX135357 |
| Anti-SARS-CoV-2 ORF7a | GeneTex | GTX632602 |
| Anti-SARS-CoV-2 spike | GeneTex | GTX632604 |
| Anti-SARS-CoV-2 Membrane | GeneTex | GTX636245 |
| Anti-SARS-CoV-2 ORF9b | GeneTex | GTX136053 |
| Anti-SARS-CoV-2 NSP5 | ProteinTech | 29286-1-AP |
| Anti-SARS-CoV-2 NSP8 | GeneTex | GTX632696 |
| Anti-ATG7 | Cell Signaling | 8558 |
| Anti-ATG9A | Abcam | ab108338 |
| Anti-OPTN | Santa Cruz | sc-166576 |
| Anti-ACE2 | R & D systems | AF933 |
| Anti-Tubulin | Sigma Aldrich | T6199 |
| Anti-Actin | Santa Cruz | sc-47778 |
| Anti-p-IRF3 | Cell Signaling | 4947 |
| Anti-IRF3 | Cell Signaling | 11904 |
| Anti-p-STAT1 | Cell Signaling | 9167 |
| Anti-STAT1 | Cell Signaling | 14994 |
| Anti-MDA5 | Cell Signaling | 5321 |
| Anti-RIG-I | Cell Signaling | 3743 |
| Anti-TRIM22 | Invitrogen | PA5-51964 |
| Anti-p-TBK1 | Cell Signaling | 5483 |
| Anti-TBK1 | Cell Signaling | 3504 |
| Anti-STING | R & D systems | MAB7169 |
| Anti-MAVS | Santa Cruz | sc-166583 |
| Anti-TOMM20 | Abcam | ab56783 |
| Donkey anti-Rabbit IgG HRP Linked Whole Ab | Sigma Aldrich | GENA934-1ML |
| Donkey anti-Mouse IgG HRP Linked Whole Ab | Sigma Aldrich | GENA931-1ML |
| Mouse anti-goat IgG-HRP | Santa Cruz | sc-2354 |
| Goat anti-Rabbit IgG (H+L) Secondary Antibody, Alexa Fluor 488 | Thermo Scientific | A-11034 |
| Goat anti-Human IgG (H+L) Cross-Adsorbed Secondary Antibody, Alexa Fluor 594 | Thermo Scientific | A-11014 |
| Goat anti-Mouse IgG (H+L) Secondary Antibody, Alexa Fluor® 488 conjugate | Life Technologies | A-11029 |
| Goat anti-Mouse IgG (H+L) Highly Cross-Adsorbed Secondary Antibody, Alexa Fluor 594 | Life Technologies | A-11032 |
| Bacterial and virus strains | | |
| SARS-CoV-2 WA1 | BEI | NR-52281 |

| | | |
|---|---|---|
| SARS-CoV-2 Alpha | BEI | NR-54971 |
| SARS-CoV-2 Omicron BA.1 | BEI | NR-56461 |
| SARS-CoV-2 Omicron BA.5 | BEI | NR-58616 |
| Sendai virus Cantell | Charles River | 10100774 |
| Vesicular stomatitis virus | J. Rose | [93] |
| Chemicals, peptides, and recombinant proteins | | |
| DMSO | Sigma-Aldrich | D2650 |
| VPS34 IN1 | Selleck Chemicals | S7980 |
| Remdesivir | MedChem Express | HY-104077 |
| Ruxolitinib | Invivogen | TLRL-RUX |
| Interferon beta (IFNβ) | PeproTech | 300-02BC |
| BafilomycinA | Sigma-Aldrich | B1793 |
| H-151 | Invivogen | inh-h151 |
| Critical commercial assays | | |
| BCA protein assay | Thermo Scientific | PI-23225 |
| DNase I | Zymo Research | E1010 |
| RNA Clean and Concentrator-25 kit | Zymo Research | R1018 |
| SYBR green master mix | Thermo Scientific | 4368708 |
| Moloney murine leukemia virus (M-MLV) reverse transcriptase | Thermo Scientific | 28025021 |
| Experimental models: Cell lines | | |
| Calu-3 | ATCC | HTB-55 |
| VERO E6 | ATCC | CRL-1586 |
| VERO-TMPRSS2 | Matt Friedman | [94,95] |
| AIR-100 | MatTek Corporation | AIR-100 |
| iAT2 | iPSC Core at UPENN | N/A |
| Oligonucleotides, primers and siRNAs | | |
| The sequences are listed in Supplemental information Table 1, and 2. | This paper, IDT, Sigma Aldrich | N/A |
| Software and algorithms | | |
| ImageJ | ImageJ.org. | N/A |
| Prism (V9.4.1) | GraphPad | N/A |
| MetaXpress 5.3.3 | Molecular Devices | N/A |
| Biorender | Biorender.com | N/A |

**Supplemental table 2. The list of qPCR primers used in this study.**

| Name | Forward (5` - 3`) | Reverse (5` - 3`) |
|---|---|---|
| SARS-CoV-2 N (V1) | ATGCTGCAATCGTGCTACAA | CCTCTGCTCCCTTCTGCGTA |
| SARS-CoV-2 nsp14 | TGGGGYTTTACRGGTAACCT | AACRCGCTTAACAAAGCACTC |
| 18s rRNA | AACCCGTTGAACCCCATT | CCATCCAATCGGTAGTAGCG |
| IFNB | CAACTTGCTTGGATTCCTACAAAG | TATTCAAGCCTCCCATTCAATTG |
| IFNL1 | CACATTGGCAGGTTCAAATCTC | AAGCCTCAGGTCCCAATTC |
| TRIM22 | CTGTCCTGTGTGTCAGACCAG | TGTGGGCTCATCTTGACCTCT |
| IFIT1 | CAACCAAGCAAATGTGAGGA | AGGGGAAGCAAAGAAAATGG |
| MAVS | CAGGCCGAGCCTATCATCTG | GGGCTTTGAGCTAGTTGGCA |
| MDA5 | TCGAATGGGTATTCCACAGACG | GTGGCGACTGTCCTCTGAA |
| STING1 | CACTTGGATGCTTGCCCTC | GCCACGTTGAAATTCCCTTTTT |
| RIG-I | CTGGACCCTACCTACATCCTG | GGCATCCAAAAAGCCACGG |
| ATG7 | ATGATCCCTGTAACTTAGCCCA | CACGGAAGCAAACAACTTCAAC |
| ATG9A | TGTTTCTCAATGAATGGAGCCTC | AAGTTAGCGATGCCAATCCAC |
| OPTN | CCAAACCTGGACACGTTTACC | CCTCAAATCTCCCTTTCATGGC |
| ACE2 | ACAGTCCACACTTGCCCAAAT | TGAGAGCACTGAAGACCCATT |
| PIK3C3 | CCTGGAAGACCCAATGTTGAAG | CGGGACCATACACATCCCAT |
| SeV | GGAGGAATCTAGGATCATACGAGGC | GCGGTAAGTGTAGCCGAAGCCG |
| VSV | CGGAGGATTGACTAATGC | ACCATCCGAGCCATTCGA |
| SARS-CoV-2 sgRNA-F | ACAAACCAACCAACTTTCGA | |
| sgRNA-ORF3a-R | | TCCTTGATTTCACCTTGCTTC |
| sgRNA-ORF6-R | | ACTGTATGCAGCAAAACCTG |
| sgRNA-N-R | | GAATCTGAGGGTCCACCAAA |
| 5’UTR-R | | GAGTTACTCGTGTCCTGTC |

Supplemental table **3. The list of siRNAs used in this study.**

| Name | siRNA sequence (5` - 3`) |
|---|---|
| ACE2 siRNA-1 | CUUUGUCACUGCACCUAAATT |
| ACE2 siRNA-2 | CAUCGAUAUUAGCAAAGGATT |
| ACE2 siRNA-3 | GGAUCCUUAUGUGCACAAATT |
| TMPRSS2 siRNA-1 | CAAUGUCGAUAUCUAUAAATT |
| TMPRSS2 siRNA-2 | CCAUGGAUACCAACCGGAATT |
| ATG2A siRNA-1 | ACAAGUUCCUGUACCUACATT |
| ATG2A siRNA-2 | CGACCUACAUGGUAUCUAUTT |
| ATG2B siRNA-1 | CCAGCGAUGUUGUCCAUAUTT |
| ATG2B siRNA-2 | GCCCGUUAAUAAGACCGAUTT |
| ATG4A siRNA-1 | GGAUCUUAGGGAAGCAGCATT |
| ATG4A siRNA-2 | GCUGUUGUCUGAUAUAAGUTT |
| ATG4B siRNA-1 | GGAUACUGGGUAGAAAAUATT |
| ATG4B siRNA-2 | AGACUUUGGUUUACAUACATT |
| ATG5 siRNA-1 | GGAUGCAAUUGAAGCUCAUTT |
| ATG5 siRNA-2 | GAACCAUACUAUUUGCUUUTT |
| ATG7 siRNA-1 | GAAGCUCCCAAGGACAUUATT |
| ATG7 siRNA-2 | CGCUUAACAUUGGAGUUCATT |
| ATG7 siRNA-3 | GGAACACUGUAUAACACCATT |
| ATG9A siRNA-1 | GGCUUCACAUGUAUGCUCATT |
| ATG9A siRNA-2 | CCACAAACGUGAGCUGACATT |
| ATG9A siRNA-3 | GAAUAUGCAUCCACAGAGATT |
| ATG9B siRNA-1 | GGAUUACAAUGUUCUCUUUTT |
| ATG9B siRNA-2 | AUCUGUUCCUUUGCCCUUATT |
| ATG16L siRNA-1 | GGAUCCAGUUGCAAUGAUATT |
| ATG16L siRNA-2 | CAUUCGAUCAGAGAGCAUATT |
| BECN1 siRNA-1 | CAGUUACAGAUGGAGCUAATT |
| BECN1 siRNA-2 | GCAGUUCAAAGAAGAGGUUTT |
| BNIP3 siRNA-1 | GGUUUCUAUUUAUAAUGGATT |
| BNIP3 siRNA-2 | CCCAUAGCAUUGGAGAGAATT |
| BNIP3L siRNA-1 | CCAUAGCUCUCAGUCAGAATT |
| BNIP3L siRNA-2 | GAUUCUUUUGGAUGCACAATT |
| CALCOCO2 siRNA-1 | CAUUGACCUAAACAACAAATT |
| CALCOCO2 siRNA-2 | GGAGGAGCUAGAAACCCUATT |
| GABARAP siRNA-1 | AGAAGAUCCGAAAGAAAUATT |
| GABARAP siRNA-2 | CCAAAGCUCGGAUAGGAGATT |
| GABARAPL1 siRNA-1 | GAAUCCACCUGAGACCUGATT |

| | |
|---|---|
| GABARAPL1 siRNA-2 | UGAGGACAAUCAUGAGGAATT |
| GABARAPL2 siRNA-1 | CAGUUCAUGUGGAUCAUCATT |
| GABARAPL2 siRNA-2 | GCCUAACUAUGGGACAGCUTT |
| MAP1LC3A siRNA-1 | CGACCGCUGUAAGGAGGUATT |
| MAP1LC3A siRNA-2 | UGAGCGAGUUGGUCAAGAUTT |
| MAP1LC3B siRNA-1 | AUGUCCGACUUAUUCGAGATT |
| MAP1LC3B siRNA-2 | CCUUCGAACAAAGAGUAGATT |
| MAP1LC3C siRNA-1 | ACUUGCUGGUGAACAACAATT |
| MAP1LC3C siRNA-2 | AGAUCUACAGAGACUACAATT |
| NBR1 siRNA-1 | CAGAGGUCAAGGAACUUAATT |
| NBR1 siRNA-2 | GAGAACAAGUGGUUAACGATT |
| OPTN siRNA-1 | GCAUUGUCUAAAUAUAGGATT |
| OPTN siRNA-2 | GGAGACUGUUGGAAGCGAATT |
| OPTN siRNA-3 | GACUUGAAGUUGCACUCAATT |
| PIK3C3 siRNA-1 | GAGAUGUACUUGAACGUAATT |
| PIK3C3 siRNA-2 | GCAUGGAGAUGAUUUACGUTT |
| PINK1 siRNA-1 | CCUCGUUAUGAAGAACUAUTT |
| PINK1 siRNA-2 | ACAGAGACCUGAAAUCCGATT |
| PRKN siRNA-1 | CCAGUAGCUUUGCACCUGATT |
| PRKN siRNA-2 | CCAACUCCUUGAUUAAAGATT |
| RAB7A siRNA-1 | GCUGCGUUCUGGUAUUUGATT |
| RAB7A siRNA-2 | GCUAGUCACAAUGCAGAUATT |
| SNAP29 siRNA-1 | CAACCAAAGUGGACAAGUUTT |
| SNAP29 siRNA-2 | GUACUGAUGCUUACCCAAATT |
| SQSTM1 siRNA-1 | GGAGCACGGAGGGAAAAGATT |
| SQSTM1 siRNA-2 | GAUGGAGUCGGAUAACUGUTT |
| STX17 siRNA-1 | GGAGAAGAUUGACAGCAUUTT |
| STX17 siRNA-2 | CCUUUGACCAGAUCCAUGATT |
| VAMP8 siRNA-1 | GGAGUUAAGAAUAUUAUGATT |
| VAMP8 siRNA-2 | UGAAGAUGAUUGUCCUUAUTT |
| VPS39 siRNA-1 | GCAGGUCUUUGCUAAACUUTT |
| VPS39 siRNA-2 | GAGAGACUACUACCUAAUATT |
| VPS41 siRNA-1 | GAUCAGUCUUUAUGCUGAATT |
| VPS41 siRNA-2 | CUUGAGAUCUGUCAACAGATT |
| WIPI1 siRNA-1 | GCACUAUUGCUGCCCAUGATT |
| WIPI1 siRNA-2 | GAAACUCCCUGAAAACAGUTT |
| WIPI2 siRNA-1 | GCAGGUCUUCGAUACCAUUTT |
| WIPI2 siRNA-2 | GUCUGGAAACGACCAAUGATT |
